# The Human Pleiotropic Map of GWAS Associations and Therapeutic Implications

**DOI:** 10.64898/2026.04.28.721048

**Authors:** Yakov A. Tsepilov, Daniel Suveges, Daniel Considine, Szymon Szyszkowski, Xiangyu Jack Ge, Irene López Santiago, Polina Rusina, Tobi Alegbe, Vivien W. Ho, Kirill Tsukanov, Juan María Roldán-Romero, Ines A. Smit, Helena Cornu, Laura Harris, Kaur Alasoo, Alexander Predeus, Samuel Lessard, Clément Chatelain, Shameer Khader, Stephanie Yang, Anna O’Carroll, Yury S. Aulchenko, Daniel Seaton, Annalisa Buniello, Ewan Birney, Eric B. Fauman, Mark I. McCarthy, David G. Hulcoop, Gosia Trynka, Ellen M. McDonagh, David Ochoa

## Abstract

Genetic support for drug targets substantially increases clinical success rates, establishing genome-wide association studies (GWAS) as central to therapeutic hypothesis generation. However, the same genetic evidence that reveals causal gene–disease relationships simultaneously exposes organism-level safety liabilities—a dimension requiring principled, genome-wide quantification. Here we systematically analyse 100,526 GWAS to yield 789,453 credible sets and gene prioritisations for 15,641 genes, with discovery showing no saturation as GWAS expand and increase diversity. We find that 64% of GWAS-implicated genes are pleiotropic, associated with traits across multiple diseases and showing a non-linear relationship between the degree of pleiotropy and clinical success. Highly pleiotropic genes—concentrated in immune, inflammatory, and oncogenic signalling programmes—are enriched in safety-terminated clinical programmes, mouse lethal knockouts, and cancer driver genes, establishing gene-level pleiotropy as a potential measure of genetically-informed organism-level safety liability. Protein-altering variant (PAV) support amplifies therapeutic signal (OR = 6.0), yet PAV targets show higher average pleiotropy, introducing a competing safety liability. Combining PAV support with intermediate pleiotropy (2–5 therapeutic areas) resolves this tension, yielding OR = 10.3 and relative success = 4.8—a profile already satisfied by 52 approved therapies. As GWAS continue to expand in scale and resolution, these findings lay the groundwork for increasingly sophisticated target discovery strategies that yield safer and more effective therapeutic hypotheses.

## Introduction

Genome-wide association studies (GWAS) provide a systematic approach to identify causal gene–disease relationships across the spectrum of human complex traits. GWAS-supported drug mechanisms achieve a 2.6-fold greater probability of clinical success than those lacking genetic evidence ^1^. Retrospective analyses find that at least 40 germline genetic associations have been translated into approved therapies ^2^ and that approximately two-thirds of recently approved drugs currently have genetic support ^3,4^, underscoring the extent to which the genetic evidence base remains underexploited. The same genetic evidence exposes both opportunities for indication expansion and organism-level safety liabilities: drug side effects are approximately twice as likely when a target gene carries human genetic evidence for a trait similar to the adverse event ^5^.

Genetic pleiotropy—the association of a single locus with multiple phenotypically distinct traits—is the rule rather than the exception across the human genome. Empirical surveys of GWAS datasets demonstrate that approximately 90% of trait-associated loci overlap associations for multiple traits, with over half of the human genome covered by at least one reported association ^6^. This finding establishes pleiotropy as a defining feature of complex trait genetic architecture. GWAS can identify highly pleiotropic genes through context-specific non-coding variants with tissue-restricted effects, whereas burden tests lack power for highly pleiotropic genes, whose rare coding variants are depleted by purifying selection ^7^. Pleiotropy-informed methods have begun to be applied to drug target prioritisation ^8,9^, building on the principle that cross-trait GWAS evidence can reveal target biology predictive of therapeutic success, but have so far remained limited in scale or to specific disease domains. Building a comprehensive atlas of human pleiotropy to inform target selection is not, however, without challenges. The computational scale of integrating hundreds of thousands of available summary statistics ^10,11^, the predominance of European-ancestry cohorts, the requirement for high-resolution ancestry-specific fine-mapping ^12^, and the absence of a genome-wide pleiotropy-informed framework have together hindered the development of an updatable, comprehensive resource for principled target prioritisation.

Here we introduce the Open Targets Gentropy framework and report its application to 100,526 GWAS, encompassing more than one trillion variant–trait statistics. Using fine-mapping, molecular QTL colocalisation, and the Locus-to-Gene model across the full allele-frequency spectrum and a broad range of human complex traits, we identify 789,453 credible sets and causal gene prioritisations for 15,641 genes—77.9% of all protein-coding genes. We integrated these data into the Open Targets Platform to facilitate systematic drug target discovery and prioritisation ^13^. This continuously updated resource maps the phenotypic effects of human genetic variation, providing a rigorous framework to prioritise therapeutic targets based on causal evidence and comprehensive safety profiles.

## Results

### Panoramic view across 100,526 complex trait GWAS

To build the most comprehensive landscape of complex trait associations to date, we developed and applied the post-GWAS analysis framework Open Targets Gentropy (Methods) to 100,526 publicly available GWAS studies from 4,250 different publications, encompassing more than one trillion single-point statistics (Methods, Fig. 1a, Extended Data Fig. 1). The analysed GWAS included binary traits and measurements from national biobanks (e.g. UK Biobank, FinnGen, VA Million Veteran Program Biobank, BioBank Japan, China Kadoorie Biobank, deCODE Genetics Biobank, Mexico City Prospective Study and Estonian Biobank), large disease cohorts and meta-analyses (e.g. CARDIoGRAM, DIAGRAM, GIANT, INTERVAL, CHARGE, IIBDGC and PGC) (Supplementary Table 1) ^11,14,15^. Each study underwent harmonisation, quality control and manual mapping to 9,280 trait ontology terms spanning 23 therapeutic areas (TAs) ^16^. Binary traits are defined here and below as ‘diseases’, and quantitative traits other than molecular quantitative trait loci (molQTLs) as ‘measurements’. Before 2017, only 23.2% of studies included more than 10% of participants from non-European backgrounds. By the end of 2024, this proportion had increased to 35.1%, reflecting the genomic community’s efforts to enhance diversity (Fig. 1d, centre).

**Figure 1.**
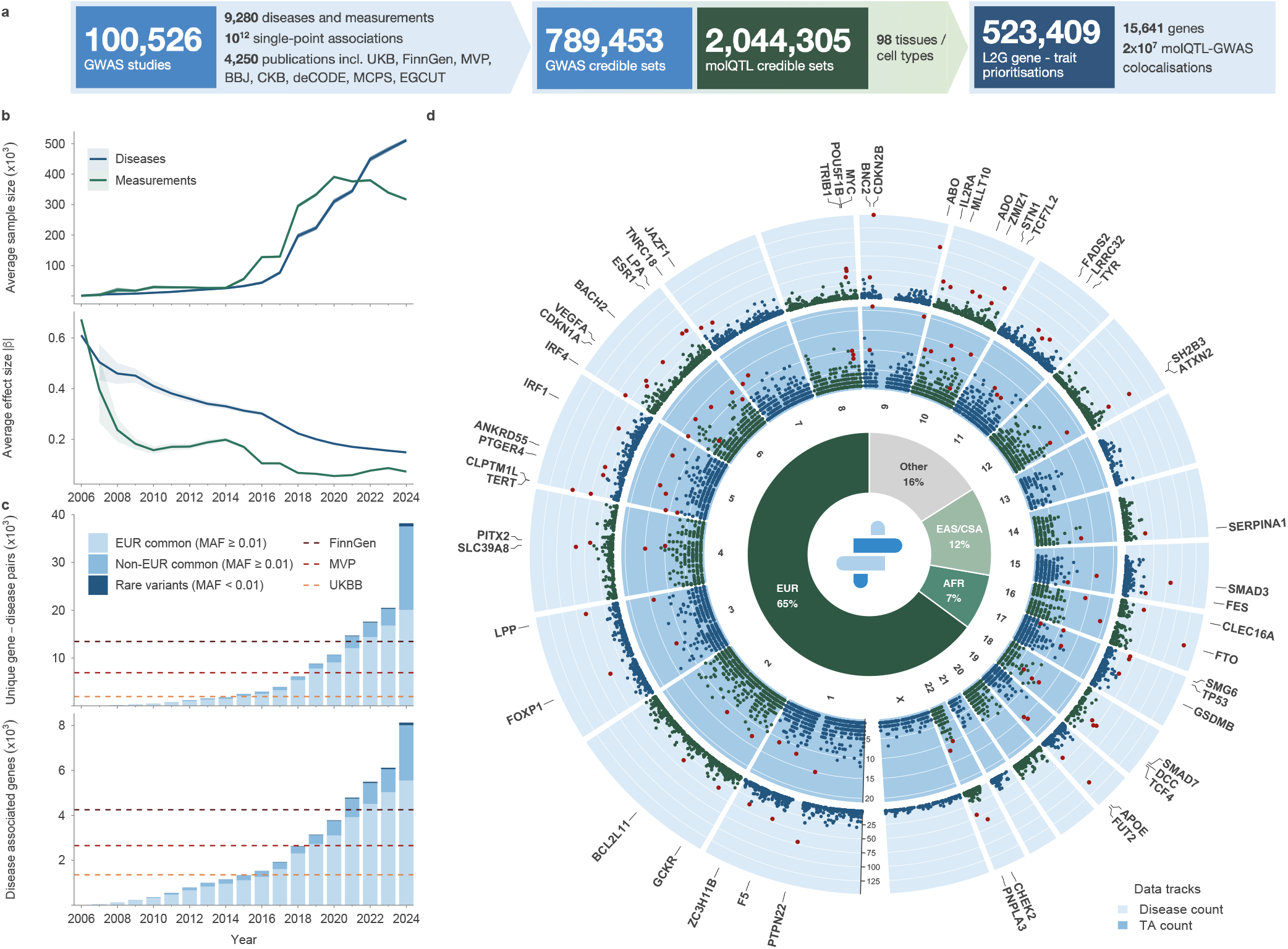
Panoramic view across 100,526 complex-trait GWAS. **(a)** Overview of study inputs and outputs. **(b)** Temporal trends in GWAS sample sizes and absolute effect sizes. Lines represent cumulative averages across all studies published in that year or earlier. Shaded bands indicate 95% confidence intervals. **(c)** Temporal trends in the discovery of gene–disease associations and unique disease-associated genes. Each bar represents the cumulative count up to that year. Dashed lines indicate the number of gene–disease pairs from representative publications of FinnGen, MVP, and UKBB. Studies with ≥ 90% non-Finnish European participants are denoted as EUR; studies that include Finnish, mixed-ancestry, or other-ancestry cohorts with *<* 90% non-Finnish European representation are denoted as non-EUR. Associations with MAF *<* 1% are considered rare and are labelled separately, regardless of study ancestry. **(d)** Pleiotropy map of disease-associated genes arranged by chromosome around the outer ring. Each dot represents a gene associated with at least one disease. The outer circle denotes the number of unique diseases associated with the gene, and the inner circle indicates the number of unique therapeutic areas linked to the gene. Red dots and labels highlight genes associated with more than 35 diseases. The centre panel illustrates the ancestral distribution of GWAS studies.

To account for linkage disequilibrium, we applied eight ancestry-specific fine-mapping strategies, which resulted in 789,453 GWAS credible sets (CSs) derived from 39,282 studies with at least one CS (Supplementary Results 1). To minimise false positive associations, we further restricted the analysis to 520,975 qualified CSs, of which 70,618 were disease CSs and 450,357 were measurement CSs (Methods). These CSs consisted of 2,024,916 unique variants, including 211,597 lead variants (49,772 with PIP ≥ 0.9). Likely causal protein-coding genes were inferred using a revised Locus-to-Gene (L2G) machine learning model (Methods) ^17^ and supported by colocalisation against 2,044,305 molQTL CSs, including transcript, protein and splice abundances across 98 tissues or cell types (Supplementary Results 2,3) ^18,19^. The analysis yielded 523,409 CSgene prioritisations, comprising 36,858 gene–disease pairs and 150,360 gene–measurement pairs (Methods). This encompassed 15,641 genes—8,285 associated with diseases and 15,160 with measurements. Altogether, these genes covered 1,394 diseases and 3,412 measurements. In total, 77.9% of all protein-coding genes were associated with at least one quantitative or binary human complex trait.

Retrospective analysis of studies conducted over the last two decades (2006–2024) revealed increasing sample sizes, enabling detection of associations with smaller mean effect sizes and a higher average number of CSs per study (Fig. 1b, Extended Data Fig. 2). Here and below, EUR denotes studies with at least 90% non-Finnish European individuals; non-EUR encompasses all others, including Finnish, mixed and other ancestry studies. Although the number of associated genes continued to increase, the rate at which genes were first discovered from disease studies based on EUR cohorts slowed (Fig. 1c). In contrast, the number of genes first associated with common variants from non-EUR studies increased in recent years. By 2024, 30% (2,469 out of 8,129) of disease-associated genes were first identified in non-EUR studies, which also contributed 16,401 of the 34,905 gene–disease associations. The contribution of rare variants to gene discovery from GWAS increased gradually, although it remains limited, partly due to our conservative quality control criteria (Supplementary Methods, Fig. 1c).

Gene-trait association discovery continued to increase, with no evidence of saturation for diseases or measurements (Fig. 1c, Extended Data Fig. 3). Gene–disease associations accumulated faster than new gene discoveries, increasing the number of diseases per gene. Out of 8,285 disease-associated genes, 5,314 were associated with two or more diseases. Similarly, 4,743 genes were associated with more than one TA, consistent with prior observations that pleiotropy is widespread across the genome (Fig. 1d).

### Selective pressures leading to concordant variant effects and directionality

Previous studies have shown that variation in complex human traits is subject to varying selective pressures, leading to distinct patterns of functional constraint ^20^. By examining 120,809 non-redundant replicated CSs with PIP ≥ 0.5 across four study types—diseases (binary traits), measurements (quantitative traits), pQTLs and eQTLs—we observed the expected inverse relationship between mean absolute effect sizes and minor allele frequency (MAF, Fig. 2a; Methods) ^21^. Lead variants with lower MAF exhibited larger effect sizes for both GWAS and molecular QTLs ^21^, reflecting in part that rare variants require larger effects to reach detection thresholds and that most studied complex traits are subject to stabilising or purifying selection ^20^. GWAS measurements exhibited the smallest effect sizes, consistent with the largest average sample sizes, followed by GWAS diseases, cis-pQTLs and eQTLs.

**Figure 2.**
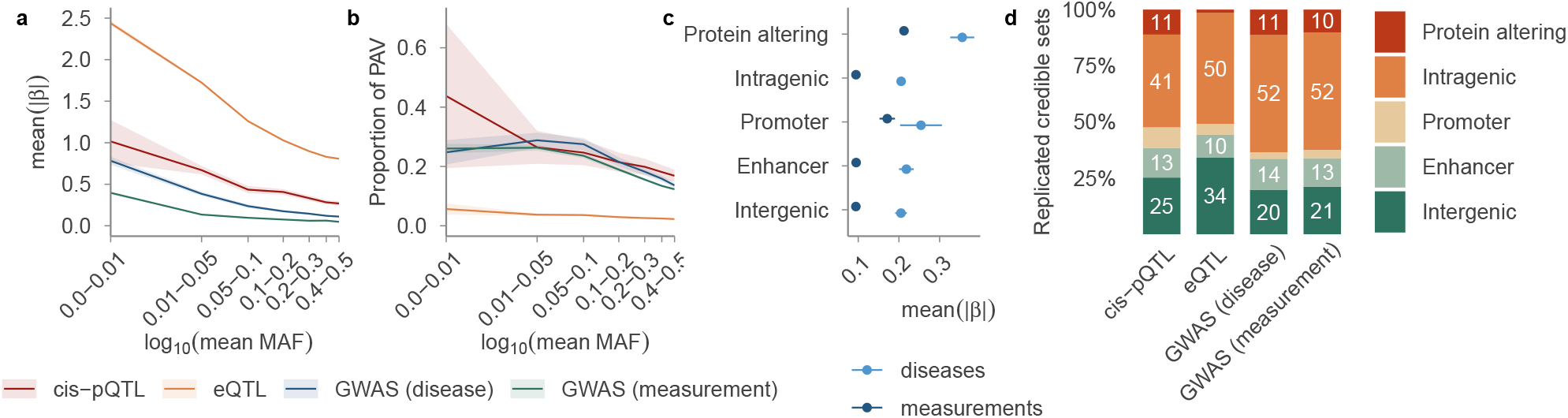
The dependency between variant effect size, MAF and predicted consequence across four study types. **(a)** Dependency between the mean absolute effect size and the MAF across four study types. Shaded bands indicate 95% confidence intervals. **(b)** The proportion of PAVs across different MAF bins. Shaded bands indicate 95% confidence intervals. **(c)** The average absolute effect size of variants across five variant consequence groups for disease and measurement GWAS. **(d)** The distribution of variant consequence/localisation across four study types. Numbers within bars indicate percentages.

Lead variants with a lower MAF in disease, measurement and cis-pQTL studies displayed a higher proportion of protein-altering variants (PAVs), a pattern not observed in eQTL CSs (Fig. 2b). PAVs displayed larger absolute effect sizes across all study types (Fig. 2c, Extended Data Fig. 4), though for eQTLs this likely reflects LD with proximal regulatory variants rather than direct coding effects on transcript abundance. Variants overlapping promoters displayed intermediate effect sizes that were significantly different from those of PAVs and other non-PAVs in eQTL and measurement studies. This difference was not significant in disease and pQTL studies, highlighting distinctive genetic constraints. Enhancer-overlapping variants were most prevalent in GWAS disease loci (13.8%), whereas promoter-overlapping variants were most prevalent in cis-pQTLs (9.1%) and least prevalent in GWAS disease loci (2.9%) ^22^. This pattern—distal enhancer enrichment in GWAS and proximal promoter enrichment in molecular QTLs—is consistent with GWAS loci acting predominantly through distal regulatory elements (Fig. 2d) ^20^. Overall, 63% of all GWAS variation fell within intragenic or PAV categories, providing a relatively straightforward basis for effector-gene assignment when variants act upon the genes in which they reside. Assigning likely causal genes to the remaining non-coding variation demands approaches beyond positional mapping alone (Fig. 2d).

### Systematic colocalisation informs non-trivial effector-gene predictions

To prioritise effector genes for GWAS signals beyond positional mapping, we enhanced the Locus-to-Gene (L2G) machine learning model ^17^. To train the L2G model, we compiled a deduplicated list of true positive effector genes (gene–trait pairs) by aggregating high-confidence associations from curated gold standards, advanced clinical trials, and extensive genetic databases (Supplementary Results 3, Methods). These pairs were then linked to the feature matrix to establish positive and negative CS-gene pairs, which were split into an 80% training and 20% held-out test set, on which the model achieved average precision of 0.81, area under the curve of 0.95 and recall of 0.65 for L2G score ≥ 0.5. We then scored all 7,066,749 CS–protein coding gene pairs and selected all those with an L2G score ≥ 0.5 as effector-gene predictions. For CSs in which no gene reached this threshold, we instead selected the highest-ranking gene if its L2G score was ≥ 0.1. The L2G model outperformed naïve prioritisation approaches (Supplementary Results 4), including nearest-gene assignment, and substantially improved upon the previous Open Targets model (Methods), achieving higher average precision (0.81 vs. 0.65) and a lower FDR (11.5% vs. 27%). As GWAS and functional genomics data accumulated over the last years, the proportion of prioritised genes lacking PAV or molQTL colocalisation support fell from 49% in 2015 to 26% in 2024, and average maximum L2G scores rose across all genes (Extended Data Fig. 6). With high-resolution fine-mapping, we found no significant differences in functional genomics feature coverage or gene predictions between primary and secondary signals (Supplementary Results 5), a critical property as GWAS sample sizes grow and secondary signals become more common.

Using eQTL colocalisation as a binary predictor of whether or not there is a colocalisation signal with an eQTL CS yielded low sensitivity (21.8%) and a high false discovery rate (FDR, 65.1%), demonstrating that this signal alone is insufficient for reliable effector-gene prediction. Of the 523,409 CS-gene assignments prioritised by L2G, 81.2% identified the nearest gene to the transcription start site. Of these nearest-gene assignments, 46.1% had no PAV or eQTL/pQTL colocalisation support (Extended Data Fig. 5). PAVs in the CS supported 13.0% of assignments, while eQTL and pQTL colocalisations supported 36.7% and 5.7% of assignments, respectively. Non-nearest gene prioritisations, though less common, often reflected strong alternative evidence. For instance, L2G prioritised *MST1* (L2G = 0.86) for an inflammatory bowel disease credible set at 3:49676792:T/C, despite being the third-closest protein-coding gene at the locus. eQTL, pQTL and splicing/transcript usage QTL (s/tuQTL) colocalisations ^23^ supported this assignment, consistent with MST1’s established role in intestinal macrophage function ^24^.

Interestingly, genes with the highest LoF constraint ^25^ in the neighbourhood of a CS were enriched among nominated effector-genes (*OR* = 1.9, *P <* 3 × 10^−90^), consistent with constrained genes being recurrently associated across diverse traits through context-specific regulatory variation ^7^.

### The interplay between functional effects and statistical power influences variant-level pleiotropy

Given the widespread pleiotropy observed across disease-associated genes (Fig. 1d), we next investigated the variant-level drivers of pleiotropy. To define the set of likely independent causal variants, we colocalised all 70,618 disease-associated CSs, yielding 20,041 independent clusters (Methods). Of these clusters, 5,595 (28%) had more than one CS lead variant, likely reflecting dense LD rather than distinct causal signals. Across these clusters, 6,678 (33%) were linked to multiple diseases (range 1–122, mean 2.16), and 4,536 (23%) were linked to multiple TAs (range 1–20, mean 1.40). Pleiotropy can be vertical, where variant’s observed effects on multiple traits are consistent with a shared causal chain such that the effect on one trait is mediated by its effect on another at the phenotypic level, or horizontal, where perturbation exerts effects on multiple traits through causal pathways that are not mediated by one another, within the limits of available genetic and phenotypic evidence. The number of unique diseases per cluster combines both forms, providing an upper bound on pleiotropy. Assuming TAs are predominantly independent, the number of unique TAs represents a lower bound on horizontal pleiotropy. The high Spearman correlation between the two counts (*ρ* = 0.81, *P <* 1 *×* 10^−16^) indicates that both measures reliably capture horizontal pleiotropy. Accordingly, we used the number of unique diseases per cluster as the variant pleiotropy score (vPS). For clusters with more than one lead variant, vPS reflects the union of diseases associated with all lead variants in the cluster.

vPS was non-linearly associated with MAF: both common and ultra-rare variants exhibited higher pleiotropy than intermediate-MAF variants (Fig. 3a). Univariate and joint linear models revealed several functional factors positively associated with vPS: GERP constraint, PAV effects, MAF, maximum effective sample size, the maximum absolute effect size (|*β*|), and predicted statistical power (Fig. 3b; Supplementary Results 6). Predicted statistical power was computed under the assumption that the variant is associated with most traits at effect sizes an order of magnitude lower than its maximal observed effect. It was jointly determined by the observed effect size (scaled down by an order of magnitude), allele frequency, and maximum sample size. In a negative binomial regression, predicted statistical power alone explained 14.7% of vPS variance, compared with 17.7% for the full joint model and only 6.0% when excluded. Maximum effective sample size acted as a technical confounder with a significant but relatively minor contribution to vPS (*R*^2^ = 0.45%).

**Figure 3.**
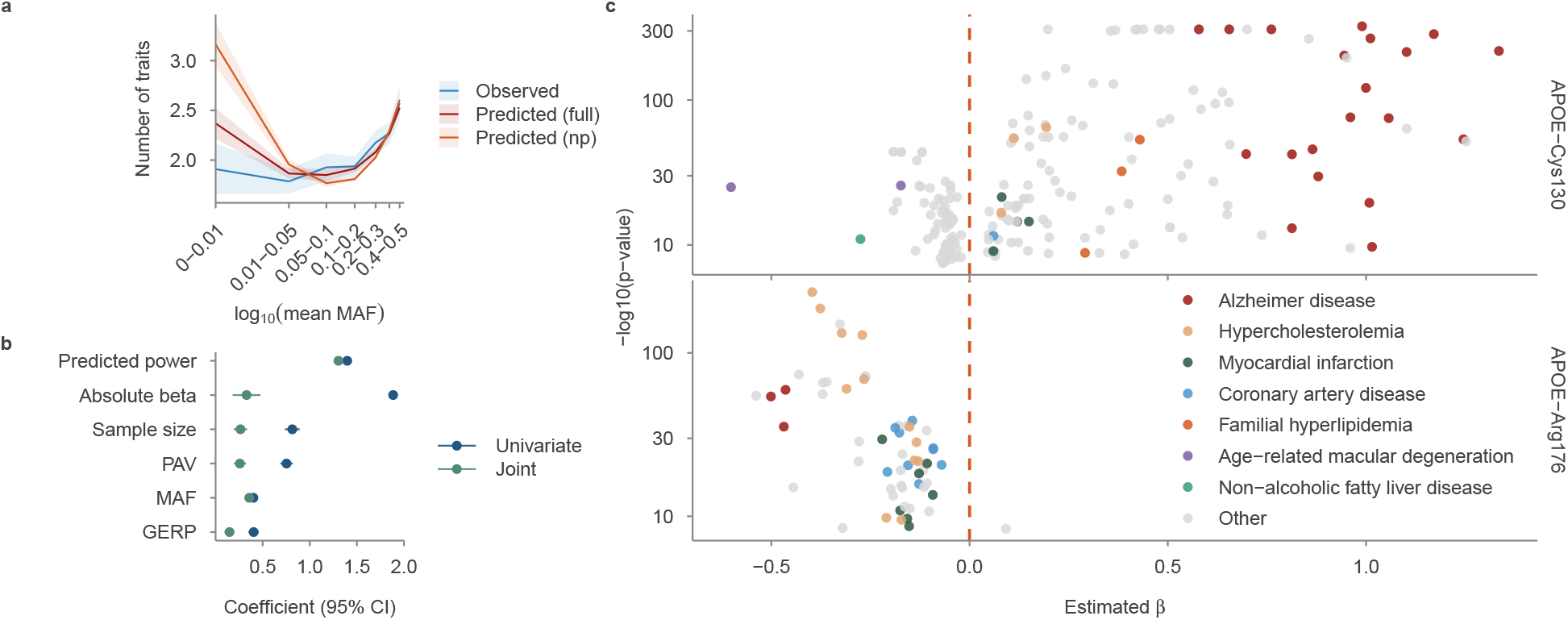
Variant-level pleiotropy modelling. **(a)** Observed and predicted variant pleiotropy scores (vPSs) in different MAF bins. The log_10_ of the mean MAF was used as the x-axis to denote each MAF bin. Two models were used to predict vPS: the full model (Predicted (full)) and the model without predicted power (Predicted (np)). The shaded band around the observed line indicates the 95% confidence interval. **(b)** The association of vPS with six covariates from negative binomial regression. Each covariate was scaled in order to have 0 as the minimum and 1 as the maximum. We used two models—univariate (each covariate separately) and joint (all covariates together). **(c)** Scatter plot of all disease associations for *APOE* variant 19_44908684_T_C (p.Cys130Arg; top) and *APOE* variant 19_44908822_C_T (p.Arg176Cys; bottom). The x-axis shows the estimated effect size (*β*); the y-axis shows − log_10_(*p*-value). Coloured points represent selected diseases; grey points represent all other associations. The orange dashed vertical line marks *β* = 0, separating associations with negative (left) and positive (right) effects.

We next investigated whether the direction of effect, increased or decreased disease risk, was consistent across all associated diseases and therapeutic areas. We defined the lead variant vPS (lead_vPS) as the number of diseases associated with a single cluster-representative lead variant. Of the 5,183 pleiotropic lead variants, 4,794 (92.5%) showed fully concordant directionality across their associated diseases, supporting previous evidence ^26^; the remaining 389 (7.5%) displayed at least one opposing direction, with discrepancies more common at higher MAFs (Supplementary Results 6). Of 135 lead variants with lead_vPS ≥ 10, 31 variants (23%) associated with 34 genes showed directionality agreements of less than 80% (Methods), demonstrating that discordant pleiotropy at the variant level is moderately present across the genome (Supplementary Table 2). Several known examples of discordant pleiotropy were identified, including variants in *APOE, ABO*, and *GCKR* ^27–29^. Variant 19_44908684_T_C in *APOE ε*4 was the most pleiotropic lead variant (lead_vPS = 85; *β* concordance = 0.66), showing associations with 15 therapeutic areas in both directions: nine with predominantly negative effects (decreased risk) and six with positive effects (increased risk) (Fig. 3c). For example, it increased the risk of Alzheimer’s disease (*β* = 1.3) and familial hyperlipidemia (*β* = 0.4), while showing protective associations with age-related macular degeneration (*β* = −0.6) and non-alcoholic fatty liver disease (*β* = −0.3). In contrast, variant 19_44908822_C_T did not exhibit discordant pleiotropy but was negatively associated with Alzheimer’s disease (Fig. 3c, bottom). Together, these findings highlight the complex landscape of variant-level associations that cannot be captured by gene-level analyses alone.

### Gene pleiotropy reflects functional specialisation and organism-level essentiality

Gene-level pleiotropy captures the tolerated range of modulation for a target, providing a framework relevant to therapeutic translation. To quantify this, we defined the gene-level pleiotropy score (gPS)—the number of unique diseases associated with all variants sharing a likely causal gene—as an estimate of apparent pleiotropy ^30^. As with vPS, gPS combines horizontal and vertical pleiotropy and is inflated by genetic correlations among diseases, making it an upper bound on pleiotropy. The number of unique TAs per gene, assuming TA independence, provides a lower bound on horizontal pleiotropy. Among the 8,285 genes associated with disease, 5,314 (64%) had a gPS greater than 1 (mean 4.45, max 148), and 4,743 genes (57%) were linked to multiple TAs (mean 2.53, max 21) (Fig. 1d). Both estimates showed a strong correlation across all genes (Spearman’s *ρ* = 0.92). The most pleiotropic genes included *FTO* (gPS = 126), *APOE* (gPS = 107), and *ABO* (gPS = 105). The tumour suppressor gene *CDKN2B* exhibited the highest gPS in the genome at 148, driven by associations across 21 TAs, despite being predicted to be under low constraint (gnomAD pLI = 5.57 *×* 10^−4^, o/e = 1.15).

Retrospective analyses of pleiotropy showed a linear increase in vPS and an exponential growth in gPS over time, matched by an exponential rise in the number of coding and non-coding disease-associated variants sharing a likely causal gene (Fig. 4a). Two drivers underlay this growth: previously identified variants continued to accumulate new disease associations, and new variants were continuously linked to genes already associated with disease, often with more conclusive effects than the variants originally reported at those genes (Extended Data Fig. 6). The number of variants per gene strongly predicted gPS (Spearman’s *ρ* = 0.87), consistent with individual variants acting on specific diseases while gene-level modulation affects a broader range of diseases ^31^.

**Figure 4.**
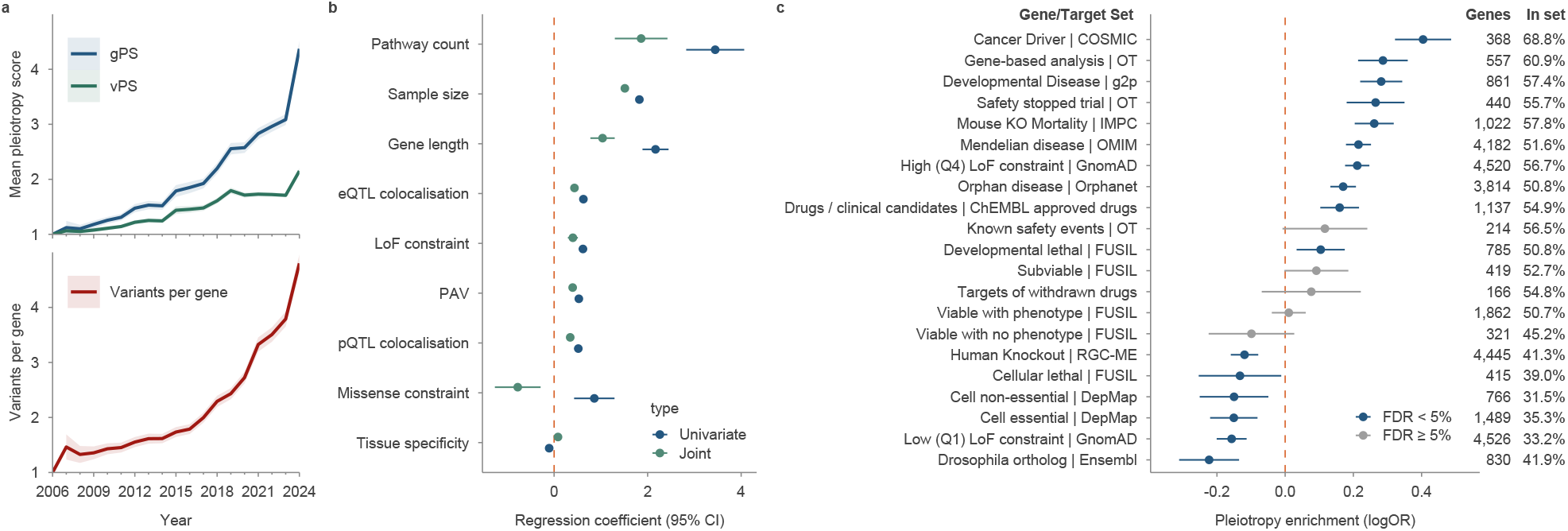
Gene-level pleiotropy modelling. **(a)** Temporal dynamics of pleiotropy levels. The top panel shows temporal trends in gene pleiotropy score (gPS) and variant pleiotropy score (vPS). The bottom panel shows the average number of unique lead variants assigned to each gene using the L2G framework; the shaded band indicates the 95% confidence interval. Each data point for a given year represents the value calculated using all studies published in that year or earlier. **(b)** Modelling of gPS using negative binomial regression. Each covariate was analysed separately (univariate model) and jointly with other covariates (joint multivariable model). **(c)** Association between log_2_-transformed gPS and membership in 21 gene sets. Each point represents the log-odds estimate from logistic regression testing the association between log_2_(gPS) and membership in a given gene set, representing the change in log-odds per doubling of gPS. Numbers to the right of each category indicate the total number of genes in the category and the percentage overlapping with GWAS genes. Blue points denote associations significant after multiple testing correction (FDR *<* 5%); grey points are non-significant.

A joint model of 9 covariates explained only 15% of gPS variance (Pearson *R*^2^ = 0.15), indicating that much of gene-level pleiotropy remained biologically unexplained. We confirmed previous observations ^32^ that gene-level pleiotropy was positively associated with several factors, including maximum sample size (univariate *β* = 1.82, *P <* 10^−308^), loss-of-function constraint (*β* = 0.64, *P <* 3.44*×*10^−40^), missense constraint (*β* = 0.59, *P <* 2.29 *×* 10^−4^), the number of pathways in which a gene participates (*β* = 3.45, *P <* 2.53 *×* 10^−16^), gene length (*β* = 2.17, *P <* 8.9 *×* 10^−53^), and the presence of PAV associations (*β* = 0.53, *P <* 6.02 *×* 10^−86^) (Fig. 4b). In the joint model, the missense constraint coefficient reversed sign, possibly reflecting collinearity with LoF constraint (Pearson *r* = 0.628). In contrast, gPS exhibited a negative correlation with the tissue specificity of the gene transcriptional profile (*β* = −0.11, *P <* 1.37 *×* 10^−3^; Methods).

To investigate the biological processes underlying pleiotropic genes, we performed gene set enrichment analysis (GSEA) using Reactome and KEGG (Methods), with the number of TAs as a quantitative measure of pleiotropy. We identified 312 significantly enriched pathways (FDR *<* 0.05; Supplementary Table 3). Highly pleiotropic genes were enriched in 221 pathways, predominantly spanning immune and inflammatory signalling, oncogenic signal transduction, and transcriptional regulation. The strongest enrichment was observed for cytokine and interferon signalling, T- and B-cell receptor pathways, JAK-STAT, NF-*κ*B, MAPK, PI3K-AKT, WNT, TGF-*β*, and receptor tyrosine kinase cascades. Collectively, these pathways define a tightly interconnected network linking immune regulation, growth factor signalling, and tumorigenesis. In contrast, low-pleiotropy genes were enriched in 91 pathways primarily representing core housekeeping functions, including DNA replication and repair, cell-cycle control, chromatin regulation, RNA processing, translation, and mitochondrial oxidative phosphorylation. This was consistent with the lower frequency of PAVs observed in these genes, as strong purifying selection limited detectable coding variation in core biological processes ^7^. Together, these findings show that pleiotropy increases in parallel with a shift from specialised genome maintenance programmes toward broad regulatory and immuno-oncogenic signalling networks.

To better understand the functional implications of gene pleiotropy, we analysed the association between log_2_-transformed gPS and membership in 21 gene sets representing varying degrees of constraint and functional roles (Methods, Fig. 4c). Using logistic regression, we evaluated whether genes with higher gPS were more likely to belong to each gene set. Higher gPS was significantly associated with membership in 10 gene sets, particularly those representing severe phenotypic consequences in multicellular organisms. These gene sets included cancer driver genes (log(OR) = 0.40, *P* = 5.6*×*10^−22^), homologs of mouse knockout-lethal genes (log(OR) = 0.26, *P* = 3.3 × 10^−19^), and developmental disorder panel genes (log(OR) = 0.28, *P* = 3.1 × 10^−19^). Drug targets involved in clinical trials terminated due to safety concerns ^33^ showed significant enrichment of higher gPS (log(OR) = 0.27, *P* = 8.9 *×* 10^−10^), demonstrating that gPS flags potential safety risks for target modulation in humans. This enrichment reflected the breadth of phenotypic involvement rather than discordant directionality: gPS retained its predictive effect on trial safety after conditioning on gene-level directionality discordance (Supplementary Results 7). Similar but non-significant trends were observed for targets of withdrawn drugs (log(OR) = 0.077, *P* = 0.30, *n* = 166) and known safety event targets (log(OR) = 0.12, *P* = 0.066, *n* = 214), likely reflecting smaller gene set sizes (Supplementary Table 4). Loss-of-function–constrained genes (Q4) derived from natural populations also showed moderate enrichment of high-gPS genes (log(OR) = 0.21, *P* = 2.1 *×* 10^−31^), whereas low-constrained genes (Q1; log(OR) = −0.16, *P* = 2.3 *×* 10^−12^) and human knockouts ^34^ (log(OR) = −0.12, *P* = 5.9 *×* 10^−9^) were enriched for lower gPS. In cellular models, the pattern reversed: high-gPS genes were depleted from DepMap essential and non-essential gene sets and FUSIL ^35^ cellular lethal genes (all log(OR) *<* 0), establishing that the organism-level signal captured by gPS was not recapitulated by cell-culture data.

### Large molecular effects with moderate pleiotropy lead to higher therapeutic success

Genetic evidence linking an on-target molecule to its indication was repeatedly associated with greater clinical trial success. We identified CSs supporting 242 approved target–indication pairs (Supplementary Table 5), a substantial increase (28%) from the 189 pairs recently reported by Minikel et al. in a smaller set of studies ^1^. Over time, the availability of GWAS evidence retrospectively validating currently approved therapies has increased, with no sign that this trend is slowing (Fig. 5a). GWAS support for approved therapies showed higher enrichment than previously reported, with *OR* = 3.62 (*P* = 4.5 *×* 10^−49^) and a relative success of *RS* = 2.76 (*P* = 3 *×* 10^−77^) (Methods, Supplementary Table 6). The enrichment pattern was largely consistent across Tas and target classes, though enzymes exhibited higher enrichment (Extended Data Fig. 7). To account for underlying heterogeneity across TAs and potential inflation driven by sample size, we applied a mixed-effects logistic regression. This adjustment reduced the overall OR from 3.62 to 3.14, confirming that both factors independently contribute to the baseline estimate (Supplementary Results 8). This estimate remained stable over time: associations with smaller effects from more recent and larger studies do not hinder drug-target identification (Fig. 5a). For example, coding loss-of-function variants in *PCSK9* showed large effects on LDL cholesterol and coronary heart disease risk ^36^, while an independent common upstream variant (rs11206510) was associated with coronary artery disease with a modest effect (|*β*| = 0.068) ^37^. Both classes of evidence correctly identified one of the most successful drug targets of the past decade, illustrating that even marginal-effect associations can reliably nominate highly successful targets.

**Figure 5.**
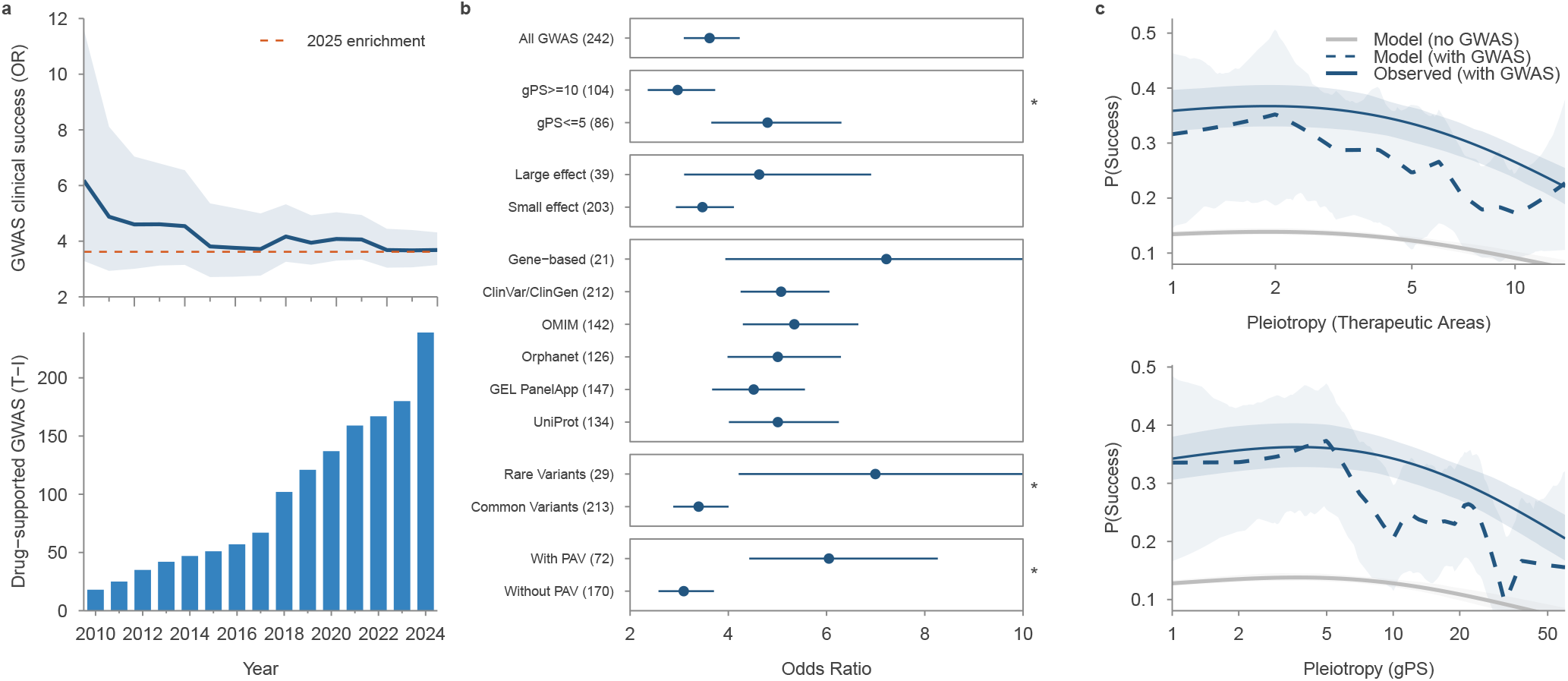
Genetic evidence and therapeutic implications. **(a)** Temporal dynamics of drug–target enrichment. The top plot shows the odds ratio (OR) of enrichment as a blue line with 95% confidence intervals shown as light blue shading; the orange dashed line indicates the 2025 enrichment estimate. The bottom plot shows a bar chart of the cumulative number of target–indication (T–I) pairs with an approved drug by year. **(b)** Forest plot of OR (95% CI) for enrichment of genetic support in approved drug targets (Phase IV) relative to all clinical candidates (Phases I–III), across gene categories. Numbers in parentheses indicate the number of T–I pairs with an approved drug. Categories are grouped by pleiotropy (gPS), effect size, rare disease resource, variant frequency, and protein-altering variant (PAV) status. Asterisks denote statistically significant differences between pairs within each group. **(c)** Predicted logistic regression probabilities and an empirical locally weighted scatterplot smoothing (LOWESS) curve. The top panel shows enrichment across the number of therapeutic areas (TAs), and the bottom panel across gPS. Each plot includes the logistic regression prediction with 95% confidence band (light blue shading) for the corresponding gene group. The x-axis is on a log scale.

We further stratified the GWAS evidence to assess the therapeutic relevance of different classes of genetic support. Using logistic regression, we found that rare-variant GWAS associations (MAF *<* 0.01) showed significantly larger enrichment than common-variant associations (*OR* = 7.0 vs 3.4; *P* = 0.0077; FDR = 0.015; Fig. 5b, Methods). Consistent with this, T-I pairs derived from rare disease resources—Orphanet (*OR* = 5.1, *RS* = 3.5), OMIM (*OR* = 4.7, *RS* = 3.4), ClinVar/ClinGen (*OR* = 5.1, *RS* = 3.4), UniProt (*OR* = 5.0, *RS* = 3.4) and Genomics England PanelApp (*OR* = 4.5, *RS* = 3.2)—showed stronger OR/RS in line with previous evidence ^1^. These resources are enriched for rare variants and severe monogenic diseases, and are largely independent of GWAS evidence. Likewise, PAV associations demonstrated significantly higher enrichment than non-PAV associations (*OR* = 6.0 vs 3.1; *P* = 0.0002; FDR = 0.001; Fig. 5b). Gene-based analyses of collapsed coding variation showed increased clinical success (*OR* = 7.2, *RS* = 4.1), although the approved-drug count was small. Associations with large effects (|*β*| *>* 0.5) showed similar odds to those with smaller effects (*OR* = 4.6 vs 3.5, *P* = 0.19), consistent with previous findings that relative success is independent of genetic effect size ^1^.

Gene-level pleiotropy introduced a further stratifying dimension of clinical success: target–indication pairs supported by genes with gPS ≤ 5 were 1.6 times more likely to result in approved therapies than those with gPS ≥ 10 (*OR* = 4.8 vs. 3.0; *P* = 0.008; FDR = 0.01, Fig. 5b). Stratified analyses across gPS bins revealed a non-linear relationship between drug-target success and gPS or number of TAs (Fig. 5c). To quantify this trend, we modelled drug-target enrichment as a function of pleiotropy, using gPS and the number of TAs as pleiotropy estimates (Methods). A model with a non-linear logarithmic pleiotropy term fitted the data significantly better than linear or baseline models (*P <* 2 *×* 10^−13^ for gPS; *P <* 8 *×* 10^−16^ for number of TAs, Fig. 5c). The pattern was robust to subsampling and outlier removal and remained significant after excluding all targets with documented safety liabilities (Supplementary Results 9). The non-linear pattern did not replicate for variant effect sizes or MAF. Genes with intermediate pleiotropy showed the highest likelihood of approval, while those with high pleiotropy showed the poorest outcomes among genetically supported therapies. Even so, high-pleiotropy genes remained more successful than clinical candidates without any GWAS support, even for unrelated diseases (*OR* = 0.74, *P* = 1.8 *×* 10^−10^).

Pleiotropy did not correlate significantly with the number of approved indications per target. Target– indication pairs supported by genetic evidence showed a higher approval rate when the target had been previously approved for another indication (*OR* = 4.13), but this did not significantly exceed the baseline success rate (*OR* = 3.6). Phase transition probabilities differed across pleiotropy groups primarily at the Phase II–III transition, the stage with the highest attrition in drug development ^38^ (Extended Data Fig. 8, Supplementary Results 10). Other phase transitions (Phase I–II: 82.6–84.2%; Phase III–approval: 29.4–31.2%) did not differ significantly across pleiotropy groups after correction for multiple testing.

Integrating these observations yielded a stricter definition of genetic support that most strongly predicts clinical success. Specifically, we identified 2,734 gene–disease associations (Supplementary Table 7) supported by PAVs, with genetic support observed in 2 to 5 TAs. Target–indication pairs meeting this definition were ten times more likely to result in approved medicines (*OR* = 10.3; *RS* = 4.8). This stricter definition, matching 52 of 242 currently approved target–indication pairs (21.5%), significantly outperformed all-GWAS, gene-based and rare-disease nominations (*P <* 1.3 *×* 10^−7^ for the difference from all other GWAS evidence). The selected gene–disease pairs were not biased toward specific TAs or target classes (Supplementary Table 8). Together, these results nominate an optimal translational strategy: large molecular effects on the disease target, with minimal collateral phenotypic impact.

## Discussion

The systematic analysis of 100,526 GWAS studies reported here establishes the most comprehensive landscape of phenotypic consequences of human genetic variation yet assembled, implicating 15,641 unique genes—77.9% of all protein-coding genes—in at least one human complex trait. This resource enables principled, genome-wide target prioritisation at a scale previously unattainable. Yet the sheer breadth of this resource alone does not resolve the challenge of therapeutic translation: the same genetic evidence that reveals causal gene–disease relationships simultaneously exposes organism-level safety liabilities and presents varying potential for therapeutic translation depending on its underlying architecture. Specifically, PAVs and rare variants provide the strongest evidence of therapeutic utility. Gene-level pleiotropy, quantified as the gPS or as the number of associated therapeutic areas, captures a complementary and largely orthogonal dimension of target risk: genes with broad phenotypic footprints across multiple disease domains are enriched in organism-level essentiality and over-represented among safety-stopped clinical programmes—a liability signal not recapitulated by cellular perturbation data routinely used in target safety assessments. Defining the optimal translational strategy therefore requires not the maximisation of either criterion in isolation, but their principled integration: prioritising targets supported by impactful genetic variation while controlling for genetically informed safety liability.

The expanding scale and ancestral diversity of included studies demonstrate that GWAS discovery is far from saturation. EUR studies dominated early GWAS efforts, but non-EUR contribution has risen substantially over the past decade, and this shift has materially extended the reach of genetic discovery. Studies with more than 10% non-EUR ancestry participants now account for 30% of unique disease-associated genes (2,469 of 8,129) and 47% of all disease–gene associations (16,401 of 34,905), contributing novel gene–trait relationships through larger combined sample sizes, greater allelic diversity and the exposure of populations to distinct environmental pressures ^39^. Increased statistical power in larger, more diverse cohorts produces progressively smaller credible sets that resolve causal variants with greater precision, while also enabling detection of effect sizes too small to have been reliably identified in earlier studies. The GLP-1 receptor locus illustrates this point. In our analysis, only modest-effect non-coding associations at *GLP1R* from the Million Veteran Program cohort ^40^ genetically supported the development of agonists, now among the most successful drugs in metabolic medicine ^41^. Greater statistical power also expands coverage of rare variation. Rare variants can pinpoint causal genes with a precision unavailable from common-variant signals alone, or reveal molecular effects large enough to guide mechanistic inference. Retrospective analysis shows that GWAS signals explain an increasing share of approved therapies—242 approved target–indication pairs now carry genetic support, compared with 189 in the preceding analysis ^1^—with no evidence of saturation in gene–disease or gene– measurement association discovery.

The scale of this study rests on community-wide deposition of GWAS summary statistics in centralised resources such as the GWAS Catalog ^11^ and the eQTL Catalogue ^18^, alongside datasets generated through large-scale consortia including FinnGen ^14^ and the UK Biobank Pharma Proteomics Consortium ^19^. Translating this volume of genetic evidence into therapeutic insight requires a systematic, scalable framework that updates continuously. To address this, we developed the Gentropy framework, which underpins the Open Targets Platform and is updated quarterly as new studies are published. The Platform integrates variant-level annotation, fine-mapped credible sets, molecular QTL colocalisation, and the results of the L2G model, which assigns effector gene probabilities at each GWAS locus (Supplementary Results 11). Together, these components provide a view of complex disease genetics less susceptible to the ascertainment biases inherent to manual curation of the literature.

The systematic scope of this analysis involves methodological choices that introduce specific limitations. First, combining GWAS studies from diverse sources requires mapping heterogeneous phenotype definitions to a common disease ontology ^16^. This harmonisation enables cross-study comparison at scale but can occasionally be imprecise, as cohorts reporting the same diagnosis may represent subtly different disease aetiologies— a limitation we partially addressed by expert curation of ontology mappings. Second, we used a simplified population dictionary that assigned all GWAS studies to seven broad ancestry labels, and the fine-mapping procedure did not explicitly model multi-ancestry LD, instead relying on the major ancestry. Moreover, we used the ancestry labels and free text sample descriptions from the GWAS Catalog, as provided by the submitters, to infer the most appropriate out-of-sample LD panel. As a result, fine-mapping false discovery rates are likely inflated, although we mitigated this through extensive aggregation of functional evidence and by using replicated or qualified CSs. Future studies could reduce this limitation substantially if in-sample LD information or author-derived fine-mapping results were routinely deposited alongside summary statistics, especially for highly specific or underrepresented populations. Third, gene assignment for non-coding associations relies on functional genomic evidence that remains incomplete—46.1% of prioritised genes currently lack PAV or molQTL colocalisation support, and the L2G model shows a proximity preference, with 81.2% of assignments nominating the nearest gene. This likely reflects both genuine biology and residual model bias. The proportion of disease-associated genes lacking PAV or molQTL support has fallen from 49% in 2015 to 26% in 2024 (Extended Data Fig. 6), and as fine-mapping methods mature and more functional genomics data are integrated, each of these limitations will diminish.

The widespread gene-level pleiotropy observed here—64% of disease-associated genes with gPS *>* 1, and 57% linked to more than one TA—aligns with the broad distribution of heritability across the genome for complex traits ^42^. This pattern reflects the interconnected nature of biology, in which genes with specific, well-defined functions can influence multiple traits through downstream biochemical and physiological pathways. Spence et al. ^7^ showed that causal protein-altering variants in pleiotropic genes face strong purifying selection, reducing their frequency to the point that they evade detection by rare-variant burden tests. GWAS instead captures tissue- and condition-specific regulatory variation that modulates these constrained genes across multiple biological contexts while eluding the selective pressure acting on their coding sequence. This enables detection of a greater proportion of disease-associated variants that escape purifying selection, as well as characterisation of phenotypes associated with conditionally essential genes such as cancer driver genes, constrained only in specific cellular contexts. Pathway enrichment reinforces the interpretation of gPS as a measure of biological constraint: low-gPS genes predominate in highly conserved housekeeping programmes including DNA repair, cell cycle control, and translation, whereas high-gPS genes are concentrated in immune, inflammatory, and oncogenic signalling programmes, denoting context-specific specialisation and critical roles in organism-level viability. Low-gPS genes likely comprise a heterogeneous group: tissue-specific genes whose narrow expression restricts their phenotypic footprint, and broadly expressed housekeeping genes rendered largely invisible to GWAS by the strong purifying selection that constrains their coding and regulatory variation.

Although decomposition of horizontal and vertical pleiotropy components falls outside this study’s scope, the strong concordance between gPS and the number of associated therapeutic areas as independent proxies (Spearman *ρ* = 0.92) demonstrates that our conclusions are robust to the choice of pleiotropy metric. These estimates nonetheless reflect a complex interplay between genuine biological signals and statistical factors that requires careful interpretation; for example, while rare variants also exhibited elevated pleiotropy, variant-level modelling demonstrates that the apparent elevated pleiotropy of rare variants is partly attributable to detection of only large-effect associations. However, this constraint reflects biological selection acting on pleiotropic variants. Pleiotropic effects reduce fitness, driving down allele frequencies and thereby restricting detectable associations to those with the largest phenotypic impacts.

The pleiotropy captured by GWAS reflects organism-level dependencies not recapitulated in cellular essentiality screens: high-gPS genes are enriched among cancer driver genes (*OR* = 1.5), mouse lethal knockouts (*OR* = 1.3), developmental disorder panels (*OR* = 1.3), and safety-stopped clinical programmes (*OR* = 1.3), yet are systematically depleted from cell-line essentiality datasets. This organism-level safety information is invisible to cellular perturbation data considered in target safety assessments. This safety signal is not fully an artefact of discordant pleiotropy: gPS retained its predictive effect on trial safety outcomes after conditioning on gene-level directionality discordance, indicating that the breadth of phenotypic involvement—not solely the direction of individual variant effects—drives the safety association (Supplementary Results 7). Several mechanisms can explain why concordant pleiotropy still poses safety risk: stabilising selection constrains gene dosage such that both increased and decreased function tend to be deleterious ^7^, and many phenotypes—such as blood pressure and immune function—are clinically adverse at both extremes. Therapeutic perturbation of a broadly pleiotropic gene therefore risks unintended consequences regardless of effect directionality.

The systematic analysis presented here demonstrates that the translational value of GWAS-based genetic support is shaped by both the functional nature of the underlying variation and the pleiotropic context in which it arises. Genetic support for approved therapies is robust and growing: 242 approved target–indication pairs now carry GWAS support (*OR* = 3.62, *P* = 4.5 *×* 10^−49^; replication success ratio = 2.76, *P* = 3 *×* 10^−77^), compared with 189 in the preceding analysis^1^ . The enrichment is stable across approval cohorts and independent of effect size, confirming that even modest non-coding associations reliably nominate effective therapeutic targets. PAVs markedly amplify this signal: protein-altering variants achieve *OR* = 6.0 versus 3.1 for non-PAV genetic support (*P* = 0.0002), and rare variant support produces a comparable elevation (*OR* = 7.0 vs 3.4, *P* = 0.0077). Effect size magnitude per se confers no additional predictive value (*OR* = 4.6 vs 3.5, not significant), establishing that mechanistic specificity—not statistical magnitude—drives therapeutic relevance.

Pleiotropy poses an apparent contradiction: PAVs carry the strongest causal evidence but are also associated with higher gPS, the strongest safety signal. This is resolved by the intermediate gPS criterion: PAV support combined with 2–5 TAs captures mechanistic specificity while avoiding the systemic liability of highly pleiotropic targets. The two criteria contribute distinct translational advantages: PAV support reflecting mechanistic specificity relevant to efficacy, and intermediate gPS limiting systemic safety liability—together addressing the principal causes of clinical trial failure. High gPS attenuates clinical success in a significant non-linear relationship (logarithmic term *P <* 2 *×* 10^−13^), with targets at a gPS ≤ 5 achieving *OR* = 4.8 versus *OR* = 3.0 for a gPS ≥ 10 (*P* = 0.008). Even highly pleiotropic targets retain a genetic advantage over uninformed selection (*OR >* 1 versus *OR* = 0.74 for targets lacking any GWAS support)—gPS stratifies rather than invalidates genetic evidence. Integrating these criteria defines an optimal translational strategy: targets supported by a PAV with an intermediate gPS (2–5 therapeutic areas) achieve *OR* = 10.3 and *RS* = 4.8 (*P <* 1.3 *×* 10^−7^ compared to all other GWAS-supported targets). This profile is already met by 52 approved therapies, demonstrating the value of combining distinct lines of genetic evidence. For high-gPS genes whose horizontal pleiotropy arises from independent, tissue-specific non-coding regulatory variants, tissue-restricted therapeutic modalities may offer a route to selective phenotype targeting, extending the actionable space.

This landscape of phenotypic consequences of human genetic variation—spanning common and rare alleles, EUR and non-EUR ancestries, disease endpoints and molecular traits—constitutes a pivotal public resource for understanding the organism-level consequences of gene modulation and, by extension, for guiding the selection and safety evaluation of therapeutic targets. The systematic infrastructure that generates it is, by design, neither static nor complete: it will sharpen continuously with growing GWAS sample sizes, increasing ancestral diversity, finer fine-mapping resolution from more comprehensive LD reference panels, and expanding functional genomics datasets covering broader tissues, developmental stages, and cellular states. Each of these advances will reduce dependence on proximity-based gene assignment, extend the reach of molecular colocalisation, and deepen the power of gPS to capture organism-level safety signals that cell-based platforms cannot access. The translational framework established here—prioritising targets at the intersection of impactful genetic variation and controlled pleiotropy—provides a principled strategy for converting human genetic findings into successful drugs. As this resource grows, so will the resolution and confidence with which that strategy can be applied.

## Methods

### The Gentropy framework

The objective of the Open Targets Gentropy pipeline is to prioritise gene(s) in GWAS associated loci. The full details of each of the pipeline parts are given in corresponding sections in the Supplementary Methods; here we provide a summary overview.

The pipeline processes various types of association data depending on the input data format, including full summary statistics, curated associations (“top hits”), and credible sets. Studies are categorized into two groups: (1) complex traits and diseases (referred to as GWAS for simplicity) and (2) molecular QTLs (molQTLs), including eQTLs, single-cell eQTLs (sc-eQTLs), pQTLs, transcript usage QTLs (tuQTLs), and splicing QTLs (sQTLs).

Each study is processed in order to obtain CSs. For studies with summary statistics or curated associations, we performed study-specific clumping and ancestry-specific fine-mapping to infer CSs. For studies with existing credible sets, we unified them. Rigorous quality control and technical validation were applied to all studies and CSs.

For each variant in a CS, we performed in-silico effect prediction using multiple methods and assigned genes near the transcription start sites (TSS). We then conducted colocalisation analyses across credible sets, including GWAS vs. GWAS and GWAS vs. molQTLs. Using these results, we constructed a feature matrix and trained a machine learning model (locus-to-gene score model) to predict causal protein-coding genes with high confidence, enabling the prioritisation of causal genes and potential drug targets.

All methods are implemented and validated in Python, packaged in the open-source Gentropy library. Pipeline orchestration is managed via Google Airflow, leveraging PySpark, Dataproc, and Google Batch.

### Data sources and processing

We ingested the following types of association data:

- Curated GWAS associations (“top hits”) from the NHGRI-EBI GWAS Catalog (accessed June 2025) ^11^.
- Full summary statistics from GWAS Catalog and the UK Biobank Pharma Proteomics Project (UKB-PPP, Europeans only) ^19^.
- 95% credible sets from FinnGen (release 12) ^14^.
- 95% credible sets from eQTL Catalogue (release 7) ^18^.

Studies and credible sets were filtered for valid formats, and harmonised to ensure consistent formatting of alleles, variant IDs, and effect directions.

Prior to ingestion, each GWAS study was mapped to the Experimental Factor Ontology (EFO) ^16^ and an ancestry, while molQTLs were annotated by affected genes and mapped to biosample ontologies. For each of the GWAS studies we assigned whether the studied trait was quantitative or binary.

Moreover, we used the following data as reference datasets:

- LD matrices from gnomAD v2.1.1^25^ and LD matrices for UK Biobank (Pan-UKBB) ^43^ were used for fine-mapping.
- gnomAD v4.1^44^ and Ensembl VEP ^45^ were used for variant annotations.

Throughout this paper, studies with at least 90% non-Finnish European participants are referred to as EUR; all others are referred to as non-EUR. The LD reference panel labels used for fine-mapping (NFE, CSA, AFR, EAS, FIN) reflect the finer-grained ancestry definitions from gnomAD and Pan-UKBB.

Full details of data processing, harmonisation, and summary statistics quality control are provided in Supplementary Methods.

### Locus definition and fine-mapping

#### Clumping

To define independent loci, three complementary approaches were implemented:

- **Distance-based clumping**: Variants within *±*500 kb of the most significant genome-wide significant top variant were clumped into one locus. Applied for PICS fine-mapping (see below).
- **LD-based clumping**: Applied an *r* ^2^ threshold of ≥ 0.5 using ancestry-matched LD matrices from gnomAD v2.1.1.
- **Locus breaker**: This custom method defines fine-mapping regions in three steps. First, standard distance-based clumping is performed. Second, loci are identified by filtering variants with *P <* 1 *×* 10^−5^, clustering nearby SNPs (within 250 kb), and retaining clumps containing at least one genome-wide significant variant. Third, oversized loci (*>* 1.5 Mb) are split using lead SNPs from initial clumps, each forming sub-loci *±*750 kb. This ensures more uniform locus sizes and improves compatibility with SuSiE LD matrix input. Compared to window-based clumping, Locus breaker produces smaller, more tractable fine-mapping regions. It was used for SuSiE fine-mapping.

#### Genome-wide significance thresholds

To define genome-wide significant loci, we applied dataset-specific thresholds. For GWAS Catalog, we used the standard genome-wide significance threshold of *P* ≤ 1 *×* 10^−8^. For UKB-PPP, we applied a study-specific threshold of *P* ≤ 1.7*×*10^−11^. FinnGen provided its own study-specific fine-mapping protocol. For SuSiE-based CSs, an additional *p*-value threshold was applied to any CS lead variant within the genome-wide significant locus: for FinnGen, GWAS Catalog and UKB-PPP, *P* ≤ 1 *×* 10^−5^; for eQTL Catalogue, *P* ≤ 1 *×* 10^−3^.

#### Ancestry-specific fine-mapping

Two fine-mapping approaches were used depending on the availability of LD and summary statistics:

- **SuSiE-inf** ^46^: Used for studies with full summary statistics whose major ancestry was non-Finnish European (NFE), African (AFR), or Central/South Asian (CSA). SuSiE was configured to allow up to ten causal signals per locus. See Supplementary Methods for full details.
- **PICS** (Probabilistic Identification of Causal SNPs) ^47^: Used for studies lacking summary statistics, ancestry-specific information, or a study design incompatible with SuSiE fine-mapping. PICS uses information about proxies of the lead variant (*r* ^2^ ≥ 0.5) and approximates posterior inclusion probabilities accordingly. See Supplementary Methods for full details.

### Colocalisation

For colocalisation analysis we overlapped 95% credible sets that shared at least one variant. We calculated overlaps between GWAS CSs and between GWAS and molQTL CSs. Two colocalisation approaches were used:

- **eCAVIAR** ^48^: applied for all overlaps.
- **COLOC** ^49^: applied when both CSs in the overlap are from SuSiE or SuSiE-inf.

The direction of colocalisation was inferred from effect sizes. We considered the colocalisation signal significant for overlaps with COLOC *P*_*H*4_ ≥ 0.8 or eCAVIAR CLPP ≥ 0.01.

### Feature matrix generation and features description

For each gene assigned to a CS, we generated a feature matrix capturing 28 functional genomic features, categorised as either *individual gene features*—scored independently for each gene—or *neighbourhood features*, calculated by dividing the individual gene score by the maximum value of that feature across all genes within the same CS. This normalisation reflects a gene’s relative importance within its local genomic context.

The feature matrix was built in three steps. First, individual gene features were computed using feature-specific rules for assigning genes to CSs, scaled between 0 and 1 (except for ‘number of genes’), and stored in a long-format matrix. Second, neighbourhood features were calculated for each gene as: neighbourhood feature = individual feature *÷* maximum feature score across CS. If the maximum value was zero, the neighbourhood score was also set to zero. Third, the matrix was reshaped to wide format with nulls filled as zeros.

The individual gene and neighbourhood features are divided into three categories:

#### Distance features

- The physical distance from credible set variants to the TSS or footprint of genes.
- Mean distance features are calculated as a weighted sum of distance scores across all variants in the CS, weighted by PIP.
- Sentinel distance features are evaluated with respect to the lead variant in the CS.

#### Molecular QTL colocalisation features

- Includes both summary statistic-derived colocalisation evidence and LD-derived colocalisation evidence (e.g., eCAVIAR).
- For GWAS signals with multiple independent molecular trait signals, the maximum colocalisation score across estimates is used.

#### Variant pathogenicity features

- Reflect Variant Effect Predictor (VEP)-derived pathogenicity scores.
- VepMean is a weighted-by-PIP sum of VEP scores across all variants in the CS.
- VepMax is the maximum VEP score across all variants in the CS.

Detailed feature definitions and scoring formulas are provided in the Supplementary Methods.

### Dynamic training set and L2G model training

In the first step, we created the effector gene list—the list of EFO–gene pairs considered to be biologically true positive associations. This list was derived by combining several sources: (1) manually curated “gold standards” from the previous Open Targets Genetics portal ^17^ (medium and high confidence); (2) gene–EFO mappings containing drug identifiers that have passed Phase 3 or Phase 4 clinical trials from ChEMBL 35; and (3) gene–EFO mappings with an evidence association score ≥ 0.95 from ClinVar, UniProt, Gene2Phenotype, Genomics England PanelApp, ClinGen, UniProt Literature, and Orphanet from the 25.06 Open Targets Platform release. To ensure uniqueness of gene–EFO pairs, the combined list was de-duplicated.

In the second step, we linked the effector gene list to the feature matrix using EFOs and genes. For each EFO–gene pair, several corresponding CS–gene pairs may exist, forming the preliminary list of positive CS– gene pairs. We applied filtering to these pairs to form the final list of positive CS–gene pairs. In the final step, for each positive CS–gene pair, all other protein-coding genes assigned to the same CS were designated as negative CS–gene pairs. The dataset was split into training and held-out test sets based on unique positive genes (80%/20%, respectively) to support robust model development and assessment.

The L2G model is a binary classifier based on a gradient boosting algorithm implemented in the xgboost library. We implemented a robust cross-validation strategy to rigorously evaluate the L2G model, using the training and held-out test sets defined above. The training set was used to fit the model and for cross-validation; the held-out test set provided an unbiased assessment of final model performance. Additionally, we performed hyperparameter tuning based on the expected selectivity and average precision (AP) of the resulting model. The final model was applied to all protein-coding gene–CS pairs in the feature matrix. Relative to Mountjoy et al. ^17^, the training set was expanded considerably, the feature set was revised to better reflect causal gene biology with partial overlap with the original features (Supplementary Methods), and the model was re-tuned accordingly; performance on the held-out test set was improved (Supplementary Results 3,4).

### Therapeutic area assignment to studies

To assign studies to TAs, we took the disease EFOs from each study and mapped them to one of 22 TAs based on the disease ontology tree. Traits with the TA ‘measurement’ were excluded. If a disease did not map to a TA it was assigned the TA ‘other’. Where a disease could be mapped to multiple TAs, we used a hierarchy system to assign the TA most relevant for the disease (Supplementary Table 9). For example, ‘ovarian cancer’ was assigned to ‘cancer or benign tumour’ rather than ‘reproductive system or breast disease’ or ‘endocrine system disease’. For pleiotropy analyses, TAs were aggregated across studies, grouping at the variant or gene level, and the unique count of TAs was used as an ancillary quantitative measure of pleiotropy alongside the total count of unique disease EFOs.

### Annotation of CS lead variants by MAF and beta rescaling

To obtain the major-ancestry MAF for lead variants in each CS, we identified the population with the largest proportion of the relative sample size from the linked study, applying the following procedure: (1) if the sample was drawn from a single population, that population was used; (2) if the sample comprised populations with uneven proportions, the population accounting for the largest proportion of the total sample size was used; (3) if the sample comprised evenly distributed populations, the NFE population was preferred, or the first population reported by the authors. Using the gnomAD v4.1 joint allele frequency table for the corresponding major population, we annotated effect allele frequency (EAF) to 2,821,580 (99.6%) of lead variants. Annotation failures totalling 11,168 (0.4%) arose from either the lead variant being absent from the gnomAD allele frequency table, or allele frequencies not being defined for the inferred major population. MAF was then calculated as MAF = min(1 − EAF, EAF).

Due to the variety of methods and phenotype standardisations used to estimate marginal effect sizes across GWAS studies and fine-mapping approaches, we re-estimated effect sizes and standard errors using *Z* -scores. Studies were grouped into binary and quantitative traits based on the reported numbers of cases and controls. For binary traits we estimated the logistic regression 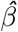, for quantitative traits we estimated the linear regression 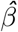. Standard errors were estimated as:

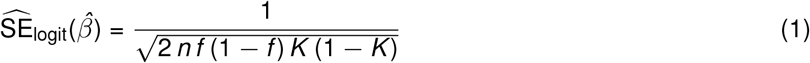

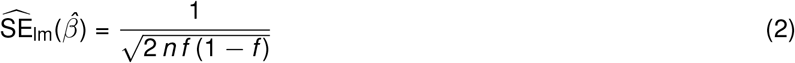

where *n* is the sample size reported by the study, *f* is the major-ancestry MAF of the lead variant, and *K* is the proportion of cases. The rescaled effect size was then estimated as:

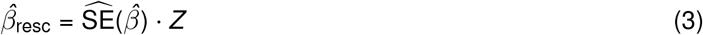

where the *Z* -score was derived from a *χ*^2^(*k* =1) distribution using the reported *p*-value from summary statistics, with the sign taken from the original study or obtained during fine-mapping. For GWAS top-hits lacking a reported 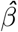, we estimated the absolute value of 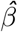 but not its direction; these variants were excluded from all analyses that require knowledge of effect direction.

### Qualified studies and CSs

Due to the systematic nature of our analysis, we expected unspecificity and noise to be present in the studies and CSs. To reduce their impact on subsequent analyses, we defined a list of qualifying studies for measurements and diseases, as well as qualifying CSs for both. The criteria for qualifying disease studies were: the trait must not be a measurement/quantitative trait, the total sample size must exceed 1,000, and prevalence must exceed 0.1%. For qualifying measurement studies we used all non-binary traits assigned as quantitative traits, excluding protein or microbiome measurements. Qualifying disease and measurement CSs were restricted to their respective qualifying study lists, then required a minor allele count of at least 20 and an absolute 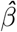 below 3. Additionally, if a CS lead variant had MAF *<* 1%, the CS was required to satisfy at least one of: a significant colocalisation with molQTLs, a PAV, or replication. In total, we identified 61,885 qualified measurement studies and 15,730 qualified disease studies, corresponding to 450,357 qualified measurement CSs and 70,618 qualified disease CSs.

### Replication of CSs

A GWAS CS was considered replicated if its lead variant was reported to be associated with the same trait (EFO term) at least twice, but in a different cohort or biobank, a different publication, or a different ancestry group.

A molQTL CS was considered replicated if its lead variant was reported to be associated with the molecular levels of the same gene at least twice.

### Assignment of most severe variant consequence

We annotated credible set variants with VEP predictions using Ensembl (release 114) annotations with canonical transcript consequences for genes located within *±*500 kb of the variant ^45^. This process generated a variant index, which was then used to annotate credible set lead variants. For each lead variant, VEP assigned the most severe consequence within the transcript range based on VEP consequence ranking ^50^.

We filtered annotated lead variants that were not replicated, did not originate from qualified CSs, or had a posterior inclusion probability below 0.5. Where a variant was reported to be associated with the same disease (or same gene for molQTLs) multiple times, we selected the single representative CS with the largest absolute effect size. Variants were then categorised according to their potential impact on protein function into PAVs and non-PAVs. Where a variant had multiple predicted impacts, the most severe was selected from the ordered list: PAV, promoter, enhancer, intragenic, intergenic.

To characterise the functional annotations of non-PAVs, we used a combination of the EPIRACTION genomic interval dataset ^22^ and VEP. EPIRACTION contains predictions from a gene-to-interval model; we filtered all associations with a score below 5%. EPIRACTION intervals are classified as promoter or enhancer regions. We overlapped lead credible set variants with these intervals to assign them to promoters or enhancers, and used VEP ranking to classify the remaining variants into intergenic and intragenic categories.

### Variant-level pleiotropy modelling

Using colocalisation, we defined clusters of colocalised disease-associated lead variants. We applied an iterative procedure starting from the most strongly associated CS and selecting all other colocalised CSs. For these CSs, we identified the set of unique lead variants and expanded the list of colocalised CSs by including all remaining CSs sharing the same lead variant. This procedure was repeated until no CSs remained unassigned. The vPS was estimated as the number of unique diseases in the cluster; for clusters with more than one lead variant, vPS was calculated as the union of diseases associated with all lead variants in the cluster.

For vPS modelling, each covariate was scaled to have a minimum of 0 and a maximum of 1. We used negative binomial regression and reported the Pearson *R*^2^ between predicted and observed values as a measure of predictive accuracy.

The expected power to detect an association was computed as:

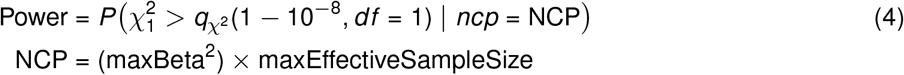

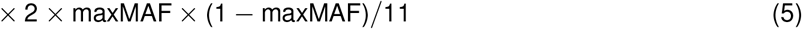

and 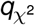 denotes the quantile of the chi-square distribution with one degree of freedom. Here, maxEffectiveSampleSize denotes the maximum effective sample size across all qualified disease credible sets reporting the variant.

Directionality of effect concordance was defined as the largest proportion of same-direction effects per variant (range 0.5–1) and was 1 for non-pleiotropic SNPs.

### Gene-level pleiotropy modelling

The gene pleiotropy score (gPS) was defined as the count of unique diseases associated with all L2G-prioritised variants sharing the same likely causal gene, with gPS = 1 indicating association with a single disease and gPS *>* 1 indicating pleiotropic association across multiple diseases.

Tissue specificity was derived from the Open Targets Platform using Human Protein Atlas (HPA) RNA expression assessments. Each gene was assigned a continuous score: tissue enriched (≥4-fold higher mRNA in one tissue vs. any other, score = 1), group enriched (≥4-fold higher in 2–5 tissues vs. any other, score = 0.75), tissue enhanced (≥4-fold higher in one tissue vs. the average of all others, score = 0.5), or low tissue specificity (score = −1). Genes with no detected expression were excluded. For modelling, tissue specificity was binarised as tissueSpecificityBinary = 1 if score *>* 0.75 (tissue enriched only) and 0 otherwise.

Each covariate was scaled to a minimum of 0 and a maximum of 1. We applied negative binomial regression and reported the Pearson *R*^2^ between predicted and observed values as a measure of predictive accuracy.

For gene set enrichment analysis (GSEA), we used gene lists ranked by *z*-transformed number of therapeutic areas values and performed enrichment analysis using blitzGSEA ^51^, which estimates *p*-values from gamma-fitted null distributions of enrichment scores using the input gene set as the background. Enrichment was conducted against curated pathway collections from Reactome v93 and KEGG v115.1.

We selected the following 21 gene categories: Cancer Driver (COSMIC) ^52^; Trial Safety ^33,53^; ChEMBL approved targets ^53^; developmental disorder panel (Gene2Phenotype) ^54^; gene-based analysis (Open Targets) ^13^; gnomAD LoF constraint Q1 and Q4^25^; mouse knockout mortality ^55^; OMIM genes ^56^; Orphanet genes ^13^; essential and non-essential genes (DepMap) ^57^; known safety targets ^13^; withdrawn drugs ^53^; human knockouts ^34^; Drosophila distant orthologs; and five categories from FUSIL ^35^. The Trial Safety gene set comprised targets of clinical trials terminated due to safety concerns, derived from ChEMBL-curated clinical trial data via the Open Targets Platform (release 25.06); oncology targets were not explicitly excluded. Each gene category was mapped to Ensembl gene identifiers, and all non-protein-coding genes were filtered out. For each category, we assessed overlap with all disease-associated genes (*N* = 8,285) and fitted a logistic regression: category membership ∼ log_2_(gPS). Statistical significance was estimated using a Wald test, followed by Benjamini–Hochberg correction for multiple testing.

### Clinical trials success modelling

We adopted the approach described in Minikel et al. ^1^ to estimate the likelihood that medicines supported by GWAS evidence—linking an on-target molecule to a disease indication—progress through the clinical development pipeline starting from Phase I. Target–indication (T–I) data were extracted from the ChEMBL evidence source in the Open Targets Platform (release 25.06) and filtered to retain only records with both target and indication information. Drugs were linked to gene targets using the manually curated mechanism-of-action assignment in ChEMBL, which reflects the target believed to be responsible for the therapeutic effect. For each T–I pair, we assigned the maximal clinical trial phase observed and included only pairs with evidence starting from Phase I.

For each genetic evidence category, we constructed a 2 *×* 2 contingency table and applied Fisher’s exact test, with columns corresponding to approval status (approved/Phase IV versus not approved/Phases I–III) and the presence or absence of genetic evidence. All oncology-related indications were excluded. The final ChEMBL dataset comprised 37,377 T–I pairs, including 6,163, 14,410, 12,240, and 4,564 pairs with maximal clinical phases I, II, III, and IV, respectively. We estimated both the odds ratio (OR) and the relative risk (denoted relative success, RS). Genetic evidence was propagated through disease ontology to capture indirect associations and was combined with ChEMBL data by requiring an exact match between disease indications. Genetic evidence categories included GWAS-based support derived from Gentropy-processed CSs (stratified by variant functional consequence, MAF, and gene-level pleiotropy), rare disease resource annotations (Orphanet, OMIM, ClinVar/ClinGen, UniProt, and Genomics England PanelApp), and gene-based rare variant associations, each assessed independently.

For stratified analyses, we used a logistic regression framework in which two independent binary predictors were defined for genetic evidence: 0 indicating absence and 1 indicating presence of genetic evidence for a given stratum. Regression coefficients were compared between models using a *t* -test. *P*-values were adjusted for multiple testing using the Benjamini–Hochberg procedure. Only ORs were estimated for this analysis, as the framework does not allow estimation of RS.

To model the non-linear relationship between clinical trial success and pleiotropy, we extended the framework by adding two covariates: the log of the number of unique diseases (or TAs) plus one, and its squared term:

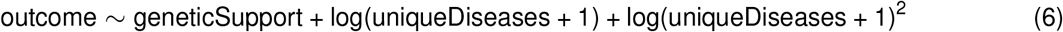

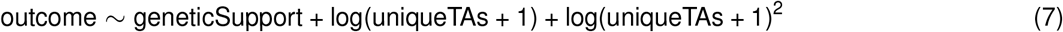

These models were compared with a baseline model (outcome ∼ geneticSupport) and with a model including only the linear term, using likelihood ratio tests. Predicted logistic regression probabilities and an empirical locally weighted scatterplot smoothing (LOWESS) curve were calculated across a grid of pleiotropy values. Ninety-five percent confidence intervals for all curves were estimated using a non-parametric bootstrap resampling approach with 200 iterations, based on the 2.5th and 97.5th percentiles of the predicted values.

## Acknowledgements

We thank Dr. Sodbo Sharapov and Dr. Nicola Pirastu for fruitful discussions on fine-mapping during the development of this work. We thank Dr. Chris Wallace for discussions about credible set colocalisation. We are also grateful to the GWAS Catalog submitters for their enormous contribution to advancing the field, and to the broader scientific community for benchmarking efforts and for providing constructive feedback on the Open Targets Platform, which has helped improve its robustness and usability. We thank all participants of cohorts underlying the GWAS data.

## Author contributions

Y.A.T. and D.O. conceived and designed the study. Y.A.T., D.C., D.Su., S.S., X.J.G., I.L.S., P.R., T.A., V.W.H., K.T., and D.O. performed the data analysis. All authors participated in the interpretation of the results, scientific discussions, and refinement of the study design at different stages of the study. G.T., E.M.M., and D.G.H. supervised the research. Y.A.T. and D.O. drafted the manuscript. All authors critically reviewed and approved the final manuscript.

## Ethics declarations

### Data availability

All results including large-scale fine-mapping are released under a CC0 licence to facilitate drug target discovery and research in statistical genetics of complex traits, and are available from https://platform.opentargets.org/downloads. The platform (https://platform.opentargets.org/) and underlying data are updated quarterly, with ongoing expansion of GWAS coverage and continuous refinement of the target-prioritisation framework.

### Code availability

The Gentropy codebase is available at https://github.com/opentargets/gentropy. The code required to reproduce the analysis described in the paper can be found at https://github.com/opentargets/Gentropy-manuscript.

## Extended Data figures

**Extended Data Fig. 1.**
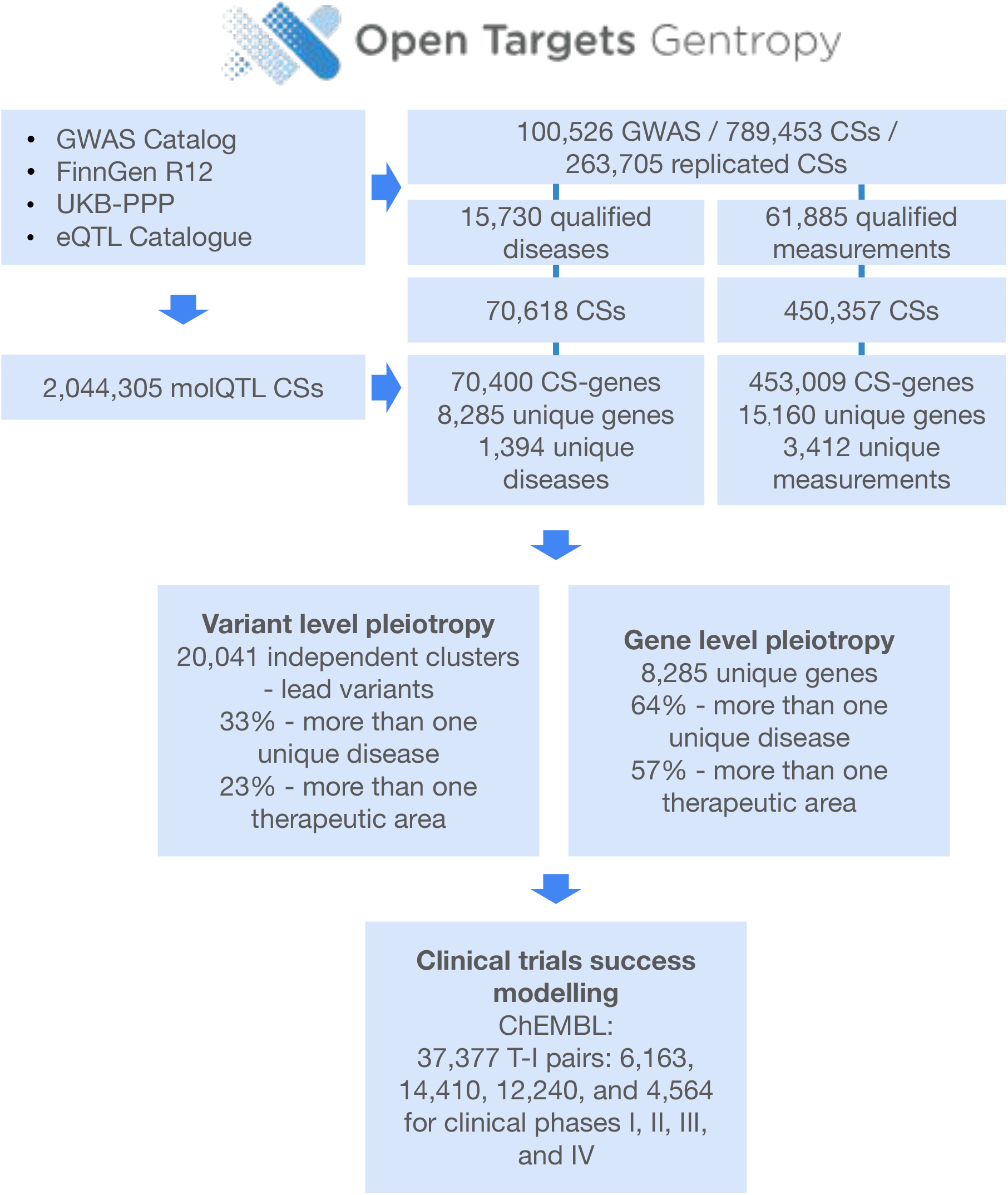
Flowchart showing the data points and results for the paper. Data flow from four input sources (GWAS Catalog, FinnGen R12, UKB-PPP, eQTL Catalogue) through fine-mapping, colocalisation, and L2G prioritisation to final outputs. Key summary statistics are shown at each stage.

**Extended Data Fig. 2.**
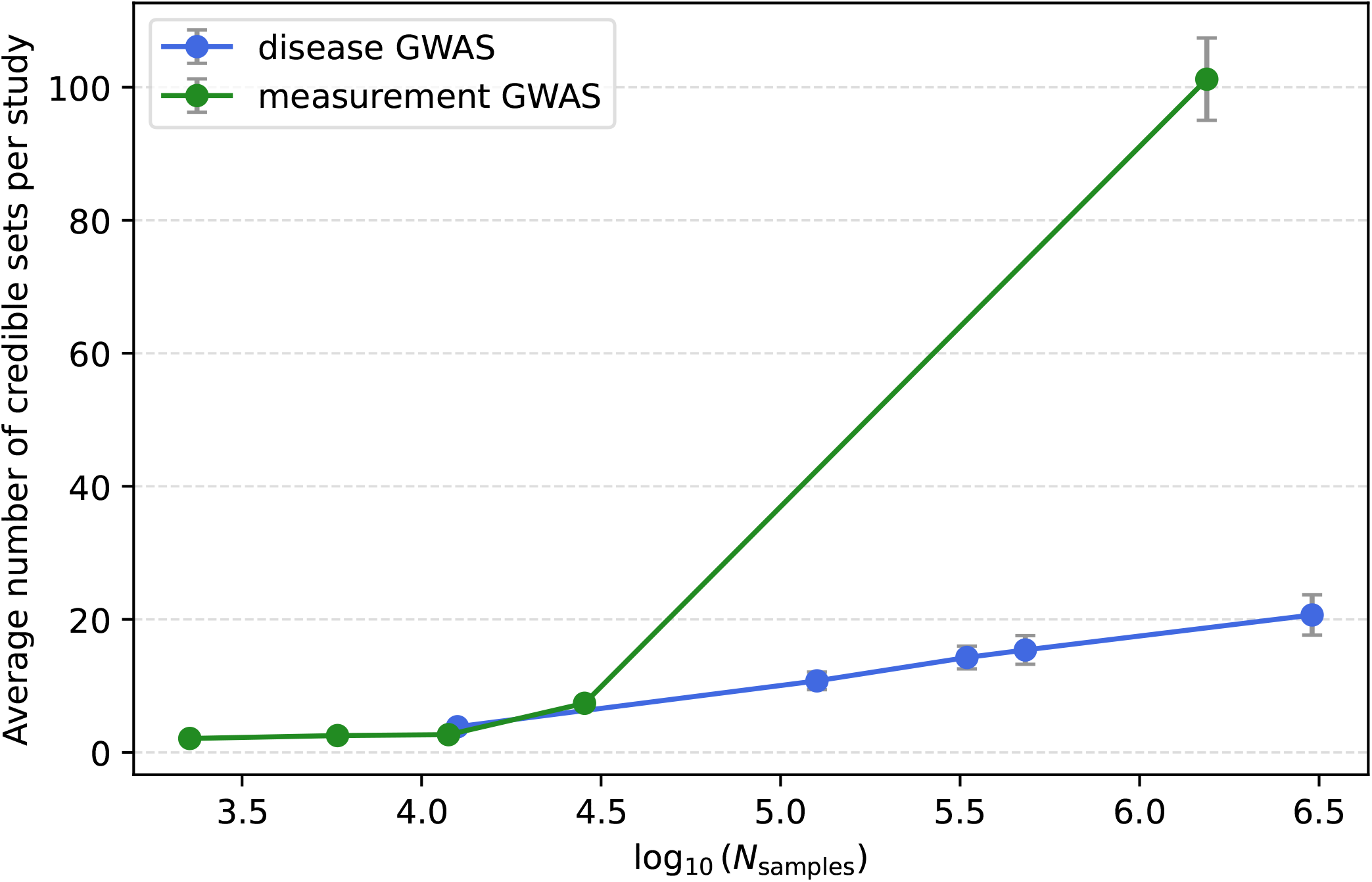
Dependence of the average number of CSs on the logarithm of GWAS sample size. The x-axis shows log_10_(*N*_samples_) bin midpoints for five quantile bins. The y-axis shows the average number of credible sets per study. Blue: disease GWAS; green: measurement GWAS.

**Extended Data Fig. 3.**
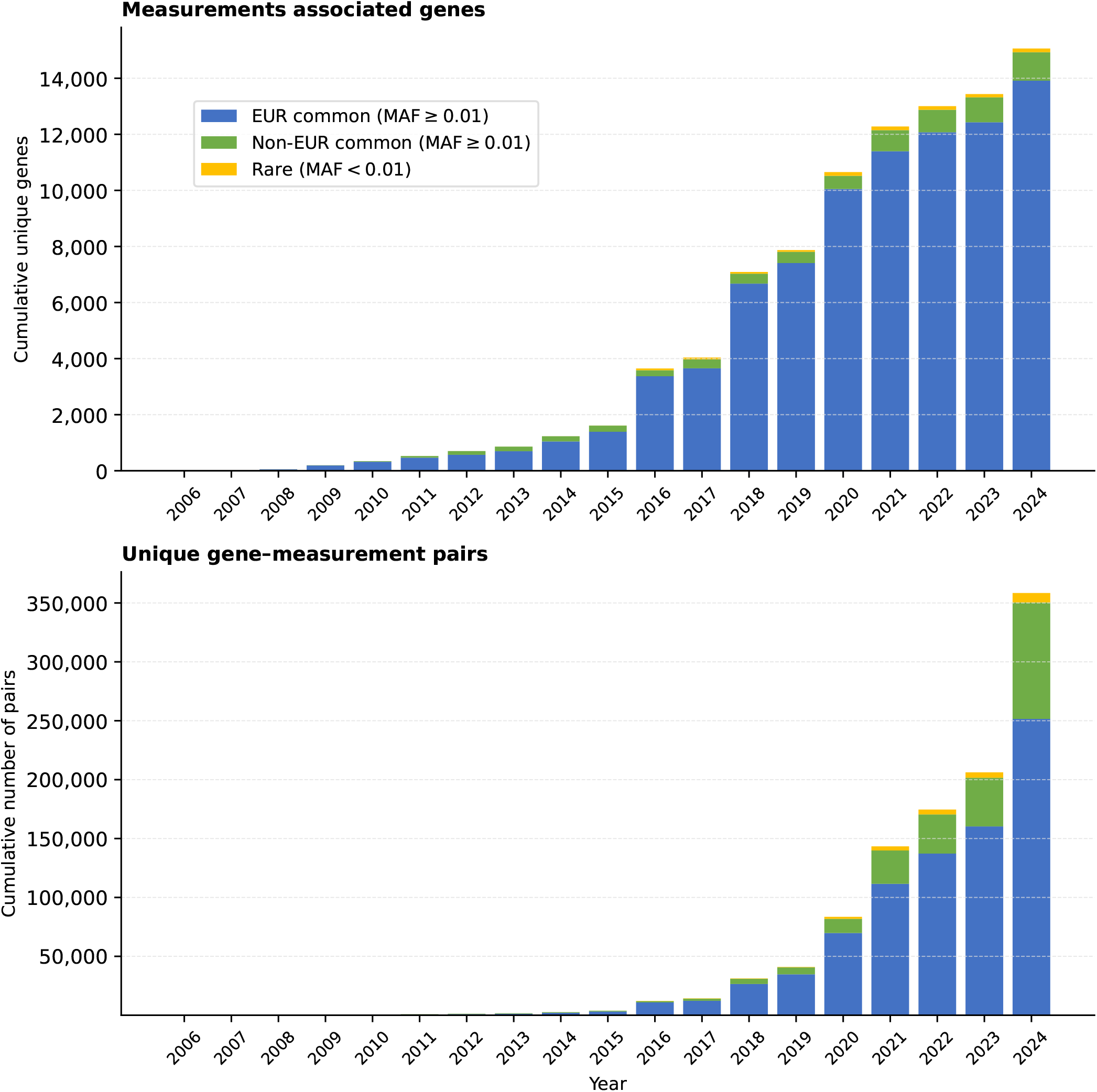
Temporal trends in the discovery of novel gene–measurement associations and unique measurement-associated genes. Top panel: cumulative number of unique measurement-associated genes by year. Bottom panel: cumulative number of unique gene–measurement pairs by year. Bars are stratified by variant frequency and ancestry: EUR common (MAF ≥ 0.01; studies with ≥ 90% non-Finnish European participants), Non-EUR common (MAF ≥ 0.01; studies with *<* 90% non-Finnish European representation, including Finnish, mixed-ancestry, and other-ancestry cohorts), and rare (MAF *<* 0.01).

**Extended Data Fig. 4.**
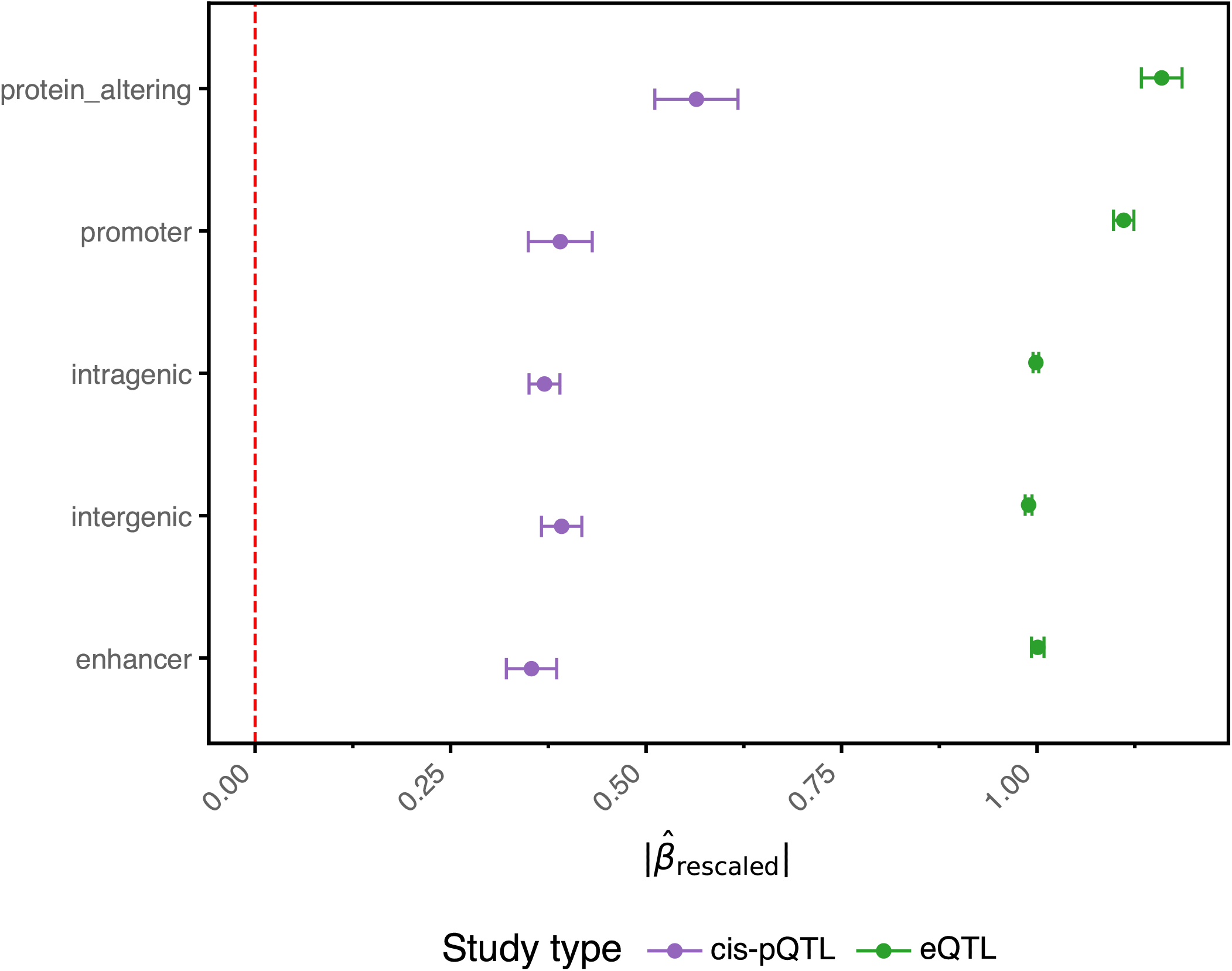
The average absolute effect size across variant consequence groups for eQTLs and cis-pQTLs. Mean absolute rescaled effect size 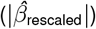 with 95% confidence intervals across five variant consequence groups for eQTLs and cis-pQTLs. The red dashed vertical line marks zero.

**Extended Data Fig. 5.**
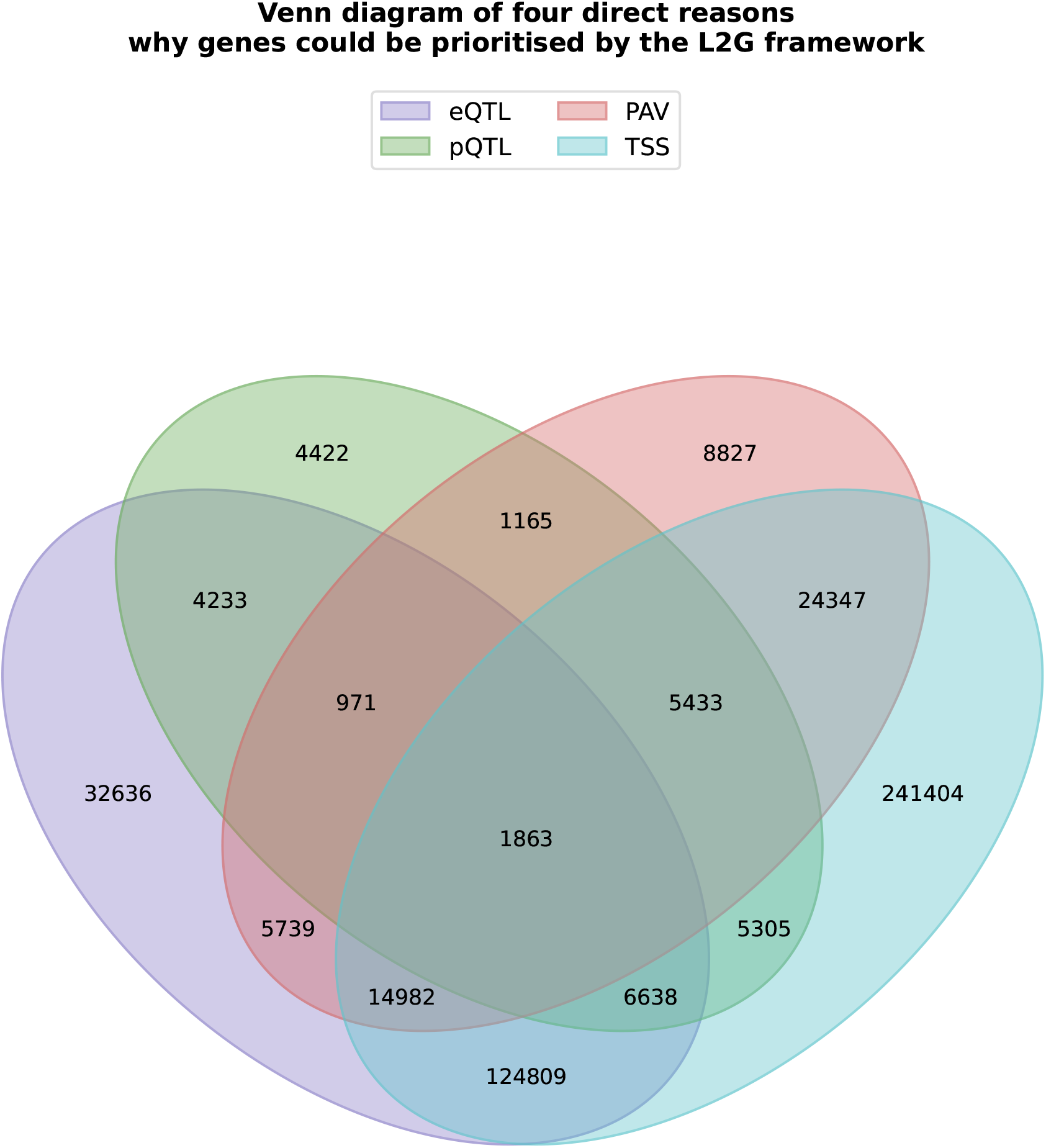
Venn diagram of four direct reasons why genes could be prioritised by the L2G framework. Four-way Venn diagram showing the overlap between four potential reasons for L2G gene prioritisation: eQTL colocalisation (the credible set colocalises with an eQTL for the gene), pQTL colocalisation (the credible set colocalises with a cis-pQTL for the gene), PAV (the credible set contains a protein-altering variant in the gene), and TSS (the gene has the nearest transcription start site to the credible set lead variant). Numbers indicate the count of CS–gene prioritisations in each overlap region.

**Extended Data Fig. 6.**
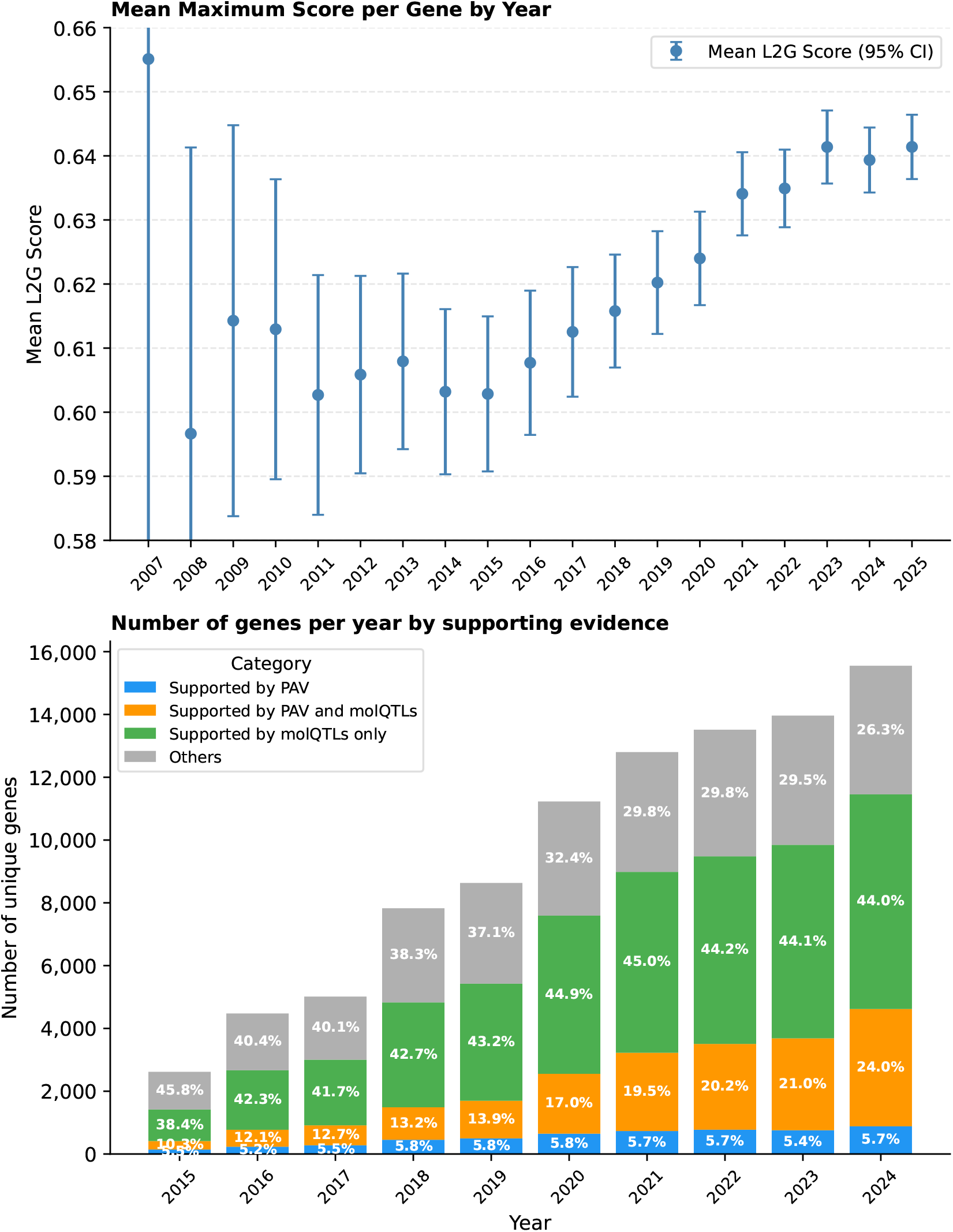
Temporal evolution of L2G confidence and supporting evidence. We examined whether gene-level L2G scores improve over time and whether the proportion of genes supported by molQTL colocalisation or protein-altering variant (PAV) evidence increases. For each year, we selected all GWAS evidence available up to that year. For each gene, we recorded the maximum L2G score and whether eQTL colocalisation or PAV evidence was present, then calculated the mean and standard error of the mean. The top panel shows the mean maximum L2G score per gene over time, with error bars indicating the standard error of the mean. The bottom panel shows the number of genes per year stratified by supporting evidence category: supported by PAV only, by PAV and molQTL colocalisation, by molQTL colocalisation only, or by neither (Others). Percentages are shown within bars.

**Extended Data Fig. 7.**
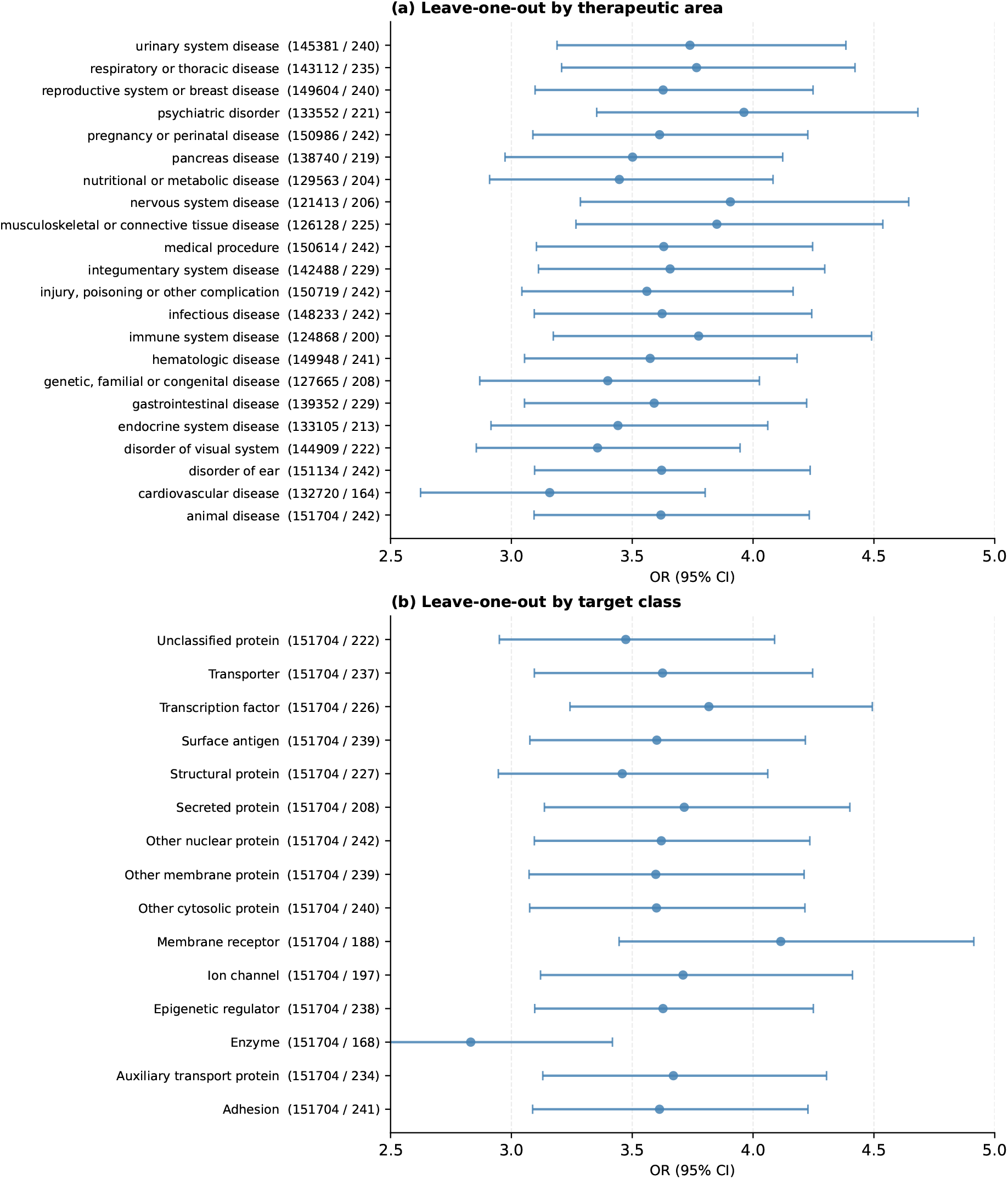
Leave-one-out analysis of successful drug target enrichment. **(a)** Leave-one-out analysis across therapeutic areas. Each row shows the OR (95% CI) for drug target enrichment obtained after excluding the indicated therapeutic area. Numbers in parentheses indicate the number of approved T–I pairs remaining after exclusion. **(b)** Leave-one-out analysis across target classes. Each row shows the OR (95% CI) obtained after excluding the indicated target class, with the same conventions as panel (a).

**Extended Data Fig. 8.**
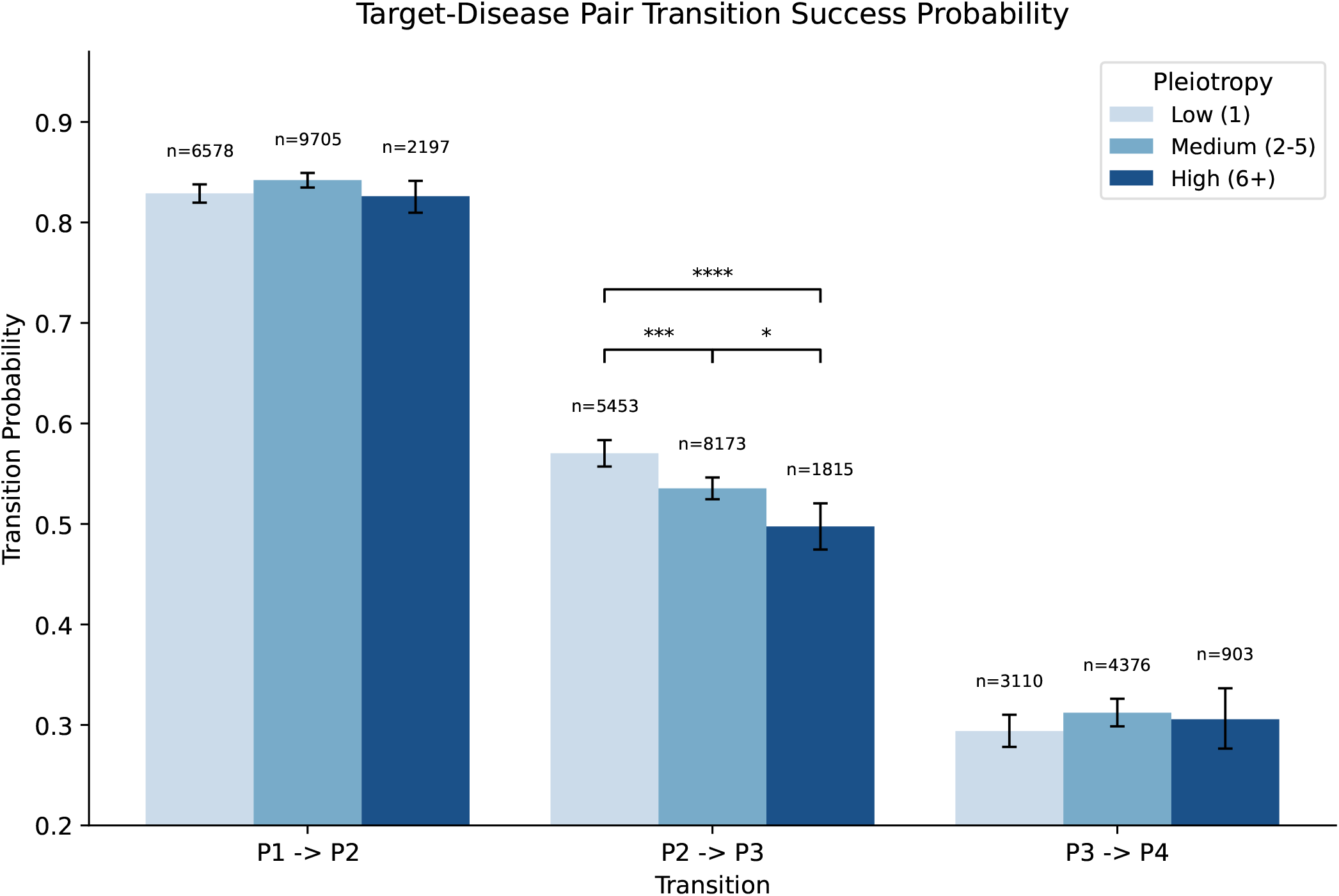
Target–disease translation success probabilities stratified by pleiotropy. Grouped bar chart of phase-to-phase transition success probabilities (Phase I → II, Phase II → III, Phase III → approval) for 18,480 target–indication pairs with GWAS genetic support, stratified by pleiotropy level: Low (1 TA), Medium (2–5 TAs), and High ( ≥ 6 TAs). Error bars show Wilson 95% confidence intervals. Sample sizes (*n*) are shown above each bar. Significance brackets above the Phase II → III bars indicate BH-FDR corrected pairwise comparisons (**** *P <* 0.0001; *** *P <* 0.001; ** *P <* 0.01; * *P <* 0.05). Phase I → II and Phase → III approval transition rates did not differ significantly across pleiotropy groups after multiple testing correction (see Supplementary Results 9).

## Supplementary information

Supplementary Methods, Results and References are available as a separate supplementary file.

**Supplementary Table 1** | **GWAS studies included in the analysis**. List of all GWAS data sources analysed, including national biobanks (UK Biobank, FinnGen R12, VA Million Veteran Program Biobank, BioBank Japan, China Kadoorie Biobank, deCODE, Mexico City Prospective Study, Estonian Biobank), large disease cohorts and meta-analyses (CARDIoGRAM, DIAGRAM, GIANT, INTERVAL, CHARGE, IIBDGC, PGC), and their key metadata (study identifier, number of samples, ancestry, trait ontology mapping, therapeutic area assignment).

**Supplementary Table 2** | **Lead variants with discordant pleiotropic effects**. The 31 lead variants (out of 135 with lead vPS ≥ 10) showing beta-effect directionality agreement *<* 0.8, with associated gene symbols, variant identifiers, number of associated diseases, number of associated therapeutic areas, directional concordance score, and selected trait associations in both directions.

**Supplementary Table 3** | **Gene set enrichment analysis results for disease-associated and pleiotropic genes**. Full results of gene set enrichment analysis (GSEA) using Reactome and KEGG gene sets (WebGestalt/blitzGSEA), with normalised enrichment score (NES), nominal *P*, and FDR-adjusted *q*-value for all 312 significantly enriched pathways (FDR *<* 0.05). Columns: pathway identifier, pathway name, database, gene set size, NES, *P*, FDR.

**Supplementary Table 4** | **Association between continuous gPS and membership in gene sets**. Log-odds estimates from logistic regression testing the association between continuous gene-level pleiotropy score (gPS) and membership in 21 gene sets representing varying degrees of constraint and functional roles. Columns: gene set name, gene set category, number of genes in the gene set, percentage overlapping with GWAS genes, log-odds estimate, standard error, *P*, FDR-adjusted *q*-value, significance after multiple testing correction.

**Supplementary Table 5** | **All target–indication pairs from ChEMBL with GWAS genetic support annotation**. All 37,377 target–indication (T–I) pairs from ChEMBL spanning Phase I to approved (Phase IV), annotated with GWAS genetic support. Includes 4,564 approved T–I pairs, of which 242 have GWAS genetic support. Columns: Ensembl target identifier, disease EFO/MONDO identifier, Open Targets indirect association score, maximum absolute rescaled effect size, minimum MAF, maximum VEP score, highest clinical phase, number of unique associated diseases, number of unique therapeutic areas, approval status (1 = approved), and genetic support indicator (1 = GWAS support present).

**Supplementary Table 6** | **Drug target enrichment analysis results**. Odds ratios (OR) and relative success (RS) estimates with 95% confidence intervals and *P*s for GWAS genetic support enrichment across approved therapies vs clinical candidates, stratified by therapeutic area and target class.

**Supplementary Table 7** | **Gene–disease associations supported by protein-altering variants with intermediate pleiotropy (2–5 therapeutic areas)**. Gene–disease associations supported by a protein-altering variant (PAV) in the credible set and with genetic support observed in 2 to 5 therapeutic areas. Columns: gene symbol, Ensembl gene identifier, disease name, EFO identifier, therapeutic area, lead variant, consequence, L2G score, number of therapeutic areas with genetic support.

**Supplementary Table 8** | **Subgroup analysis of PAV-supported gene–disease pairs by therapeutic area and target class**. Enrichment statistics (OR, RS, *P*) for PAV-supported gene–disease pairs with intermediate pleiotropy, broken down by therapeutic area and by target class (enzyme, GPCR, ion channel, kinase, nuclear receptor, other), demonstrating absence of significant bias toward specific subgroups.

**Supplementary Table 9** | **Therapeutic area assignment**. Mapping of EFO/MONDO/OTAR ontology identifiers to the 23 Open Targets therapeutic areas used throughout the analysis. Diseases are assigned to therapeutic areas by propagating EFO ontology ancestry to the root terms listed here.

**Supplementary Table 10** | **Fine-mapping statistics by data source and fine-mapping method**. Summary of credible set (CS) counts, median and mean CS size, and proportion of single-variant CSs for each combination of data source (GWAS Catalog summary statistics, FinnGen R12, eQTL Catalogue v.7, UKB-PPP) and fine-mapping method (SuSiE-inf, PICS). Includes replication rates by ancestry group.

**Supplementary Table 11** | **Colocalisation overlap statistics**. Total numbers of pairwise overlapping credible set pairs (GWAS vs GWAS; GWAS vs molQTL), counts of significant colocalisation signals by method (eCAVIAR CLPP *>* 0.01; COLOC H4 *>* 0.8), and breakdown by QTL type (eQTL, *cis*-pQTL, *trans*-pQTL).

**Supplementary Table 12** | **Performance comparison of L2G and naïve gene prioritisation methods**. Sensitivity, specificity, positive predictive value (PPV), and false discovery rate (FDR) for each prioritisation strategy (L2G thresholds 0.5, 0.8, 0.005; eQTL colocalisation; pQTL colocalisation; protein-altering variant; nearest-to-TSS; combined approach) evaluated on the held-out test set and on the combined training and test set.

