## Supplementary Materials for "The Human Pleiotropic Map of GWAS Associations and Therapeutic Implications"

[1]Open Targets, Wellcome Genome Campus, Hinxton, Cambridgeshire CB10 1SD, UK [2]Wellcome Sanger Institute, Wellcome Genome Campus, Hinxton, Cambridgeshire CB10 1SA, UK [3]European Molecular Biology Laboratory, European Bioinformatics Institute (EMBL-EBI), Wellcome Genome Campus, Hinxton, Cambridgeshire CB10 1SD, UK [4]University of Tartu, Ülikooli 18, 50090 Tartu, Estonia [5]Target, Disease & Systems Biology, Sanofi, 350 Water Street, Cambridge, MA 02141, USA [6]Merck & Co., Inc., 320 Bent Street, Cambridge, MA 02141, USA [7]GSK, Gunnels Wood Road, Stevenage, UK [8]Pfizer Research & Development, 1 Portland St, Cambridge, MA 02139, USA [9]Genentech, 1 DNA Way, South San Francisco, CA 94080, USA ✉ (Y.T.); (G.T.); (E.M.); (D.O.)

**Terms and Definitions** 4

**Supplementary Results** 5

**1 Systematic ancestry-specific fine-mapping catalogue** 5

**2 Systematic colocalisation** 6

**3 Systematic gene prioritisation using L2G model** 6

**4 Comparison of L2G and naïve gene prioritisation methods** 8

**5 Importance of the secondary fine-mapping signals for the gene prioritisation** 8

**6 Variant-level pleiotropy modelling** 9

**7 gPS predicts trial safety independently of directional discordance** 11

**8 Drug target enrichment bias due to sample size and therapeutic areas** 12

9 Non-linearity of gPS vs drug target success 12

10 Phase transition probabilities stratified by pleiotropy 13

11 Integration to Open Targets Platform 14

Supplementary Methods 16

1 Orchestration 16

2 Data sources 17

#### Terms and Definitions

**Gentropy** — Python package to facilitate the interpretation and analysis of GWAS and functional genomic studies for target identification. This package contains implementation of methods, datasets and steps used along the Gentropy pipelines.

**SummaryStatistics** — A Gentropy dataset, a GWAS object that consists of information about all SNPs in the study. Each study has a unique studyId.

**Fine-mapping** — Statistical analysis technique used to pinpoint the specific genetic variant(s) most likely responsible for a trait association identified in a GWAS. The main result is a credible set (CS) — the minimal list of variants with assigned posterior probabilities (PIPs) to be causal that form a predefined probability. We use 95% CS, which means that the expected sum of PIPs for all variants in CSs have to be within the range of [0.95,1]. Gentropy contains multiple methods implemented to perform the fine-mapping on StudyLocus datasets, these include PICS[1] and SuSIE-inf[2].

**StudyLocus** — A Gentropy dataset that represents a genetic locus derived from GWAS study. It can be the result of clumping (the procedure for defining the lead variant in the locus), fine-mapping (see above), or a GWAS Catalog curated association (in this case it consists of information about the curated lead/top variant only, for more details see below). If it is the result of fine-mapping, it will consist of information about credible sets and the lead variant in those credible sets. We will use the term 'credible set' interchangeably with StudyLocus. Each StudyLocus has a unique studyLocusId.

**StudyIndex** — A Gentropy dataset that represents information about studies from different sources. Each study has a unique studyId.

**L2G** (Locus-to-Gene) — The Machine Learning framework to predict which gene is likely causal in the region around the significant GWAS signal. Based upon the method described in the original publication[3] with some updated adaptations as explained in more detail below.

**DAG** (Directed Acyclic Graph) — Used to describe each of the Gentropy/Platform pipelines run by the Airflow orchestration layer.

**LD** — Linkage disequilibrium.

**Genetics ETL** — The part of the Gentropy pipelines responsible for the validation of studies and credible sets, form the variant index, run colocalisation, generate the L2G feature matrix and run the L2G prediction.

**Major ancestry abbreviations:** NFE — Non-Finnish European, FIN — Finnish European, AFR — African, CSA — Central and South Asian, EAS — East Asian. See more details at https://gnomad.broadinstitute.org/news/2023-11-genetic-ancestry/.

### Supplementary Results

#### 1. Systematic ancestry-specific fine-mapping catalogue

Using the framework, we processed all studies from GWAS Catalog (v. 25.06)[4] and from FinnGen R12[5] that resulted in 100,526 GWAS studies in total. Fine-mapping resulted in 789,453 credible sets (CSs) that cover 39,282 unique studies (20.24% are binary traits) and 9,280 unique EFO terms covering all 23 therapeutic areas (TAs). The quantitative traits were represented by different measurements varying from anthropometric traits like BMI or height to metabolite levels. We excluded metagenomics or proteomics measurements from consideration.

Fine-mapping of all the studies revealed that 27.3% had at least one credible set. We observed that 32% of GWAS with at least one CS have the sample proportion of non-Finnish Europeans (NFE) ancestry less than 90%. GWAS with at least one CS cover 4,250 different publications, with the earliest publication dating back to 2006. The average and median size of the GWAS CSs were 24.61 and 5, respectively and varied across data sources and fine-mapping methods (Supplementary Table 10). The proportion of CSs with single-variant resolution (defined as CSs consisting of a single variant with PIP $> 0.9$) ranged from 13.5% (FinnGen) to 40.1% (GWAS Catalog summary statistics). The higher proportion observed in the GWAS Catalog may indicate an inflated false positive rate in fine-mapping when using out-of-sample LD or rare variants.

There was a significant but modest positive correlation between CS size and the MAF of the lead variant (slope of linear regression of size vs. MAF = 14.0, $P \lll$ 1e–16), consistent with the expectation that rare variants tend to be less linked to other variants and thus form smaller CSs. Additionally, we observed a negative correlation between the sample size of the study and the size of the CSs ($P \lll$ 1e–16), which likely reflects that larger studies have greater statistical power and therefore yield more precise fine-mapping. However, it could also suggest that larger studies are more prone to statistical errors in fine-mapping, potentially leading to a higher rate of single-variant CSs.

We processed eQTL Catalogue v.7[6] and UK Biobank Pharma Proteomics Project (UKB-PPP; Europeans only)[7] to obtain molecular QTLs (molQTLs). It resulted in 2,044,305 credible sets that cover 29,342 unique genes in 98 unique tissues/cell types (1,402,222 eQTLs, 33,731 pQTLs). See Supplementary Table 10 for detailed information on the processed data and fine-mapping.

Given the systematic nature of the analysis, we expect some degree of loss of specificity in studies, metadata assignment and fine-mapping. To minimise the impact of this on subsequent analysis, we have categorised the studies and CSs into several groups. Firstly, we defined replicated CSs (Online Methods). A total of 263,705 (33.4%) GWAS and 1,461,445 (71.5%) molQTL CSs were considered as replicated. Secondly, based on GWAS metadata, we defined qualified diseases and measurements. We also defined qualified CSs for qualified studies. It should be noted that, due to the much smaller replication rate and expected larger false discovery rate (FDR) in fine-mapping, we applied additional criteria for rare CSs (to be replicated or to colocalise with molQTLs or to have a protein-altering variant). In total, 61,885 studies were considered qualified measurements, and 15,730 were considered qualified diseases. Out of the 789,453 GWAS CSs, 520,975 were considered as qualified (450,357 and 70,618 were considered qualified measurements and disease CSs, respectively). The number of qualified CSs with the lead variant being rare (MAF $<$ 1%) was 15,311 (2.94%). All downstream analyses were performed using either qualified and/or replicated CSs.

Given the systematic nature of our analysis, we emphasise the importance of rigorous quality control (QC) at every level of the data. In our case, QC was implemented at the study, variant, disease, and gene levels (Supplementary Methods). For example, at the study level, we excluded around 30% of GWAS Catalog studies prior to the fine-mapping due to different reasons. With the increasing prevalence of pleiotropy and the growing number of associated genes, such QC measures are essential to distinguish true associations from false positives.

The most challenging and computationally intensive component of the pipeline is fine-mapping. We employed two complementary approaches with different resolutions: PICS[1] and SuSiE-inf[2]. PICS provides lower-resolution fine-mapping by ignoring secondary signals, while SuSiE is considered the gold standard, albeit computationally expensive and sensitive to GWAS and LD heterogeneity[2]. To manage these demands, we used Google Airflow to orchestrate and efficiently parallelise ancestry-specific fine-mapping, enabling us to process GWAS Catalog studies in a feasible timeframe. We did not include infinitesimal effect estimation in SuSiE-inf or LD outlier detection as implemented in CARMA[8]. As a result, we expect an inflated FDR for fine-mapping, particularly in non-NFE studies where the quality of external LD reference panels is lower. This limitation was reflected in the replication rates of CSs, which were highest in NFE studies and decreased in CSA and AFR studies.

The effect estimates and directionality of lead variants associated with the same disease showed moderate concordance across disease studies and ancestries: 16% of lead variant–disease pairs were identified in more than one GWAS study, of which only 15% showed significant Cochran's heterogeneity ($P < 10^{-4}$), due to biological or technical sources of error.

#### 2. Systematic colocalisation

For colocalisation analysis we overlapped 95% credible sets that shared at least one variant. We calculated overlaps between GWAS CSs and between GWAS and molQTL CSs. The total numbers of overlaps and significant colocalisation signals of GWAS vs GWAS and GWAS vs molQTLs CSs are outlined in Supplementary Table 11. In total 67% of all overlaps lead to the significant colocalisation by eCAVIAR[9] and 79% by COLOC[10]. Out of all qualified CSs, 330,584 (63%) had a significant colocalisation with any type of molecular QTL. Excluding *trans*-pQTL colocalisations reduces this number to 302,264 (58%). Finally, 285,229 (55%) had at least one single significant colocalisation with a protein coding gene molQTL CS. These colocalisations were assigned to 14,026 unique protein coding genes.

#### 3. Systematic gene prioritisation using L2G model

For each gene assigned to a credible set, we assessed the functional genomic features and generated a feature matrix (Online Methods). The feature matrix included 513,568 qualified CSs with at least one protein coding gene assigned to it. The total number of CS-gene pairs was 7,066,749. The average number of genes assigned per CS was 13.76, the median was 10.

To train the L2G model, we developed a procedure for dynamically constructing the training set and test sets (held-out) (Online Methods). The resulting sets are constructed in a data-agnostic manner and are expected to be less biased than training sets created through manual curation. The training and test datasets consisted of 8,520 positive gene-CS pairs corresponding to 1,377 unique positive gene-EFO pairs (390 unique genes). The training set consisted of 7,386 positive and 106,973 negative CS-gene pairs. The test set consisted of 1,134 positive and 17,477 negative CS-gene pairs. The total ratio of positives to negatives was 1:14.6. The proportion of the positive gene-EFO pairs, when the positive gene's transcription start site (TSS) was the nearest to the lead variant of CS, was 56.1% (773 out of 1,377), consistent with previously reported estimates. On the held-out test set, the model achieved average precision (AP) of 0.81, area under the curve (AUC) of 0.95. Using a threshold of L2G $\geq$ 0.5 and test set, the model shows the precision of 0.885, selectivity of 0.994 and recall of

0.645, indicating a good performance. For full details of model training and performance, see Supplementary Methods. The feature importance, AUC plot and confusion matrix are depicted on Supplementary Figure 1.

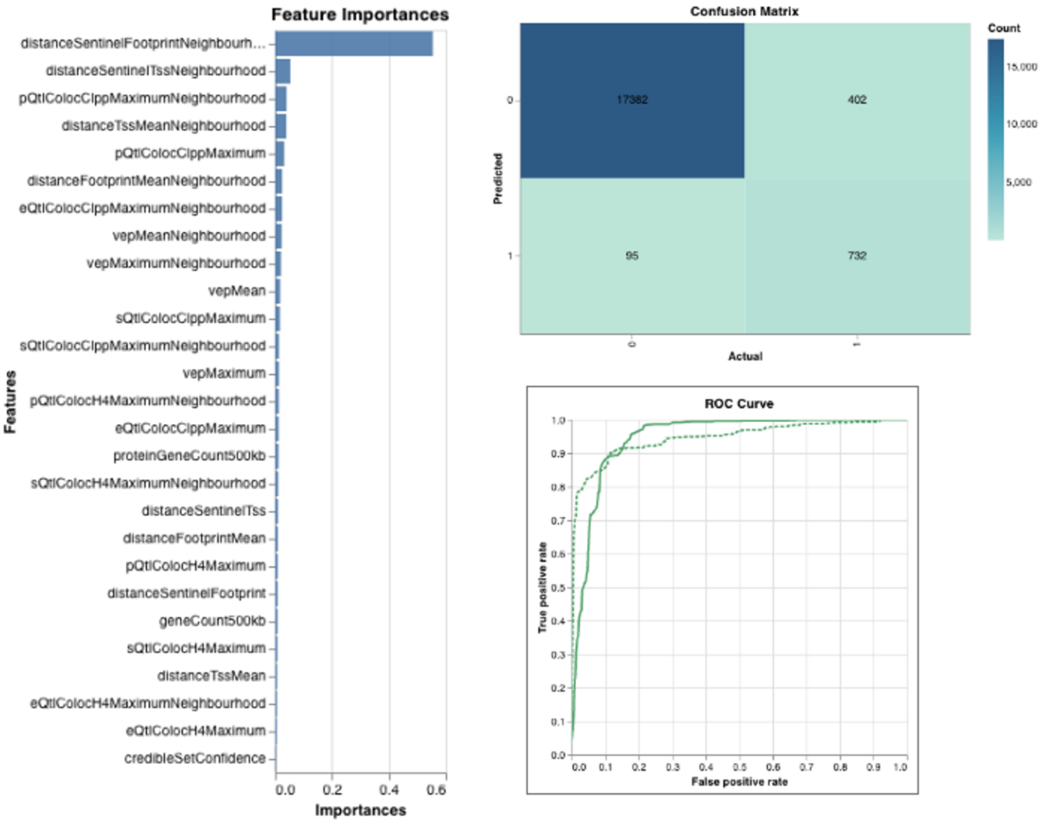

**Supplementary Figure 1.** Standard XGBoost model evaluation outputs logged via Weights & Biases (W&B) on the held-out test set. Left: feature importance scores ranked by gain (average gain contributed by each feature across all splits); higher values indicate greater contribution to predictions. Top right: confusion matrix at a classification threshold of 0.5, showing counts of true negatives (17,362), false positives (402), false negatives (95), and true positives (732), corresponding to a precision of 88.5% and a recall of 64.5%. Bottom right: ROC curve with AUC = 0.95 and average precision (AP) = 0.81.

Applied to a qualified CSs feature matrix, on average, as expected, the L2G model predicted only one gene per CS with an L2G score $\geq$ 0.5 (309,989 CSs, 59.5%). Additionally, 17,463 CSs (3.4%) had more than one gene with L2G $\geq$ 0.5, while 193,523 CSs (37.1%) had no gene with L2G $\geq$ 0.5.

We recommend using an L2G threshold of 0.5 for gene prioritisation, as it provides a reasonable balance between FDR and sensitivity. However, for 37.1% of CSs it leads to no gene assignments. For such cases, we instead use the top-ranked gene by L2G score (filtered by 0.1) as the prioritized gene. In total, using this combined approach, we prioritised 523,409 genes for 520,975 CSs comprising 15,641 unique genes (8,285 for diseases and 15,160 for measurements), covering 1,394 unique diseases and 3,412 unique measurements.

We investigated four potential reasons why the gene was associated with the trait (Extended Data Fig. 5): 36.7% (191,871) prioritised genes colocalised with the corresponding eQTLs and 5.7% (30,030) with pQTLs, 13.0% (63,327) were associated because of the protein-altering variants (PAVs) in CS, 81.2% (424,781) genes were nearest to TSS, with 46.1% (241,404) having no PAV, eQTL or pQTL evidence.

To estimate the novelty of the CSs identified, we used Open Targets Genetics data (release 22.10) as a reference. Of all GWAS CSs, 456,323 (58%) were considered novel, while 333,130 were classified as previously known.

Almost all disease-associated genes were a subset of measurement-associated genes. Interestingly, when combined with Orphanet, gene-burden associations, and eQTLs, this results in 18,950 unique genes corresponding to 94.1% of all protein-coding genes being associated with at least one human trait or disease. This is most likely an underestimation, and it is plausible that we will soon reach a point where nearly all protein-coding genes are associated with at least one trait.

#### 4. Comparison of L2G and naïve gene prioritisation methods

We compared the L2G score gene prioritisation approach using different thresholds with the combined approach used in the main text, as well as naïve gene prioritisation approaches: (1) genes whose TSS is closest to the lead variant; (2) all genes that were significantly colocalising with CSs and closest to the TSS gene; (3) all genes that were colocalising with *cis*-pQTL genes; (4) if the CS has a PAV in this gene. If the CS had a max coloc H4 $> 0.8$ or max CLPP $> 0.01$ we considered as having significant colocalisation. If it had a max variant effect predictor (VEP) score $> 0.66$ we considered it having PAV.

We evaluated FDR, recall, selectivity and precision using train+test and test-only datasets (Supplementary Materials Table 1 and Supplementary Table 12). The lowest FDRs were observed for L2G $> 0.8$ (7.7%), PAV (19.3%), combined approach (21.2%) and L2G $> 0.5$ (11.5%). The nearest-to-TSS gene approach had an FDR of 27.0%. The combined approach was better than naïve distance or molQTL prioritisation by both recall and FDR. Interestingly, eQTL colocalisation showed moderate sensitivity (21.8%) but a high FDR (65.1%), suggesting that gene prioritisation based solely on eQTL colocalisation should be approached with caution.

**Supplementary Materials Table 1.** L2G vs naïve prioritisation predictions statistics.

| | **Test dataset only** | | | | **Training and test datasets** | | | |
|---|---|---|---|---|---|---|---|---|
| **Evidence** | Sensitivity (recall) | Specificity (selectivity) | PPV (precision) | FDR | Sensitivity (recall) | Specificity (selectivity) | PPV (precision) | FDR |
| L2G $>$ 0.5 | 0.646 | 0.995 | 0.885 | 0.115 | 0.736 | 0.995 | 0.904 | 0.096 |
| L2G $>$ 0.005 | 0.839 | 0.932 | 0.443 | 0.557 | 0.880 | 0.948 | 0.538 | 0.462 |
| L2G $>$ 0.8 | 0.339 | 0.998 | 0.923 | 0.077 | 0.528 | 0.999 | 0.965 | 0.035 |
| eQTL coloc | 0.218 | 0.974 | 0.349 | 0.651 | 0.348 | 0.960 | 0.375 | 0.625 |
| pQTL coloc | 0.184 | 0.992 | 0.613 | 0.387 | 0.141 | 0.997 | 0.782 | 0.218 |
| PAV | 0.248 | 0.996 | 0.807 | 0.193 | 0.207 | 0.997 | 0.804 | 0.196 |
| Nearest | 0.702 | 0.983 | 0.730 | 0.270 | 0.644 | 0.982 | 0.708 | 0.292 |
| Combined | 0.776 | 0.986 | 0.788 | 0.212 | 0.798 | 0.989 | 0.831 | 0.169 |

#### 5. Importance of the secondary fine-mapping signals for the gene prioritisation

We used qualified diseases to perform the analysis. We used only SuSiE or SuSiE-inf based CSs. Each CS was assigned a unique study-locus pair. Each locus could have from 1 to 10 credible sets. If the locus had more than one CS we ranked them by *p*-value from lowest to highest and considered the most significant as primary signal and others as secondary. For each CS we checked the maximal value of features related to eQTL and pQTL colocalisation and VEP. If the CS had a max coloc H4 $> 0.8$ or max CLPP $> 0.01$ we considered as having significant colocalisation. If it had a max VEP score $> 0.66$ we considered it having PAV.

From 24,558 disease-associated GWAS regions, we selected those with at least one SuSiE-based CS. Of these, 6,354 regions contained both primary and secondary CSs (8,780 and 11,439 CSs, respectively). Among the 6,354 regions, 2,862 had a primary CS with no significant eQTL/pQTL colocalisations or PAV. Strikingly, 81.5% of these regions nevertheless contained a secondary CS with at least one significant functional genomic feature. The proportions of primary and secondary CSs with significant functional genomics features were broadly comparable: (1) eQTL colocalisation: 48.5% (primary) vs. 42.4% (secondary); (2) pQTL colocalisation: 9.8% vs. 7.2%; (3) PAV: 17.5% vs. 14.6%. This pattern was consistent even when the analysis was restricted to replicated CSs only.

#### 6. Variant-level pleiotropy modelling

Pleiotropy — defined as a genetic variant influencing multiple traits — was examined across disease-related traits only. We constructed a matrix of estimated effect sizes for 40,706 disease-associated variants and 1,403 diseases (each with at least one associated credible set). For disease–variant pairs reported in multiple studies, we retained the association with the largest absolute effect size. Diseases were treated as independent, and we did not distinguish between horizontal and vertical pleiotropy.

Of the 40,706 lead variants, 9,828 were associated with multiple diseases and classified as pleiotropic. As shown in Supplementary Figure 2, rare variants (MAF $<$ 1%) were underrepresented in our dataset, due to specific filtering procedures applied during quality control.

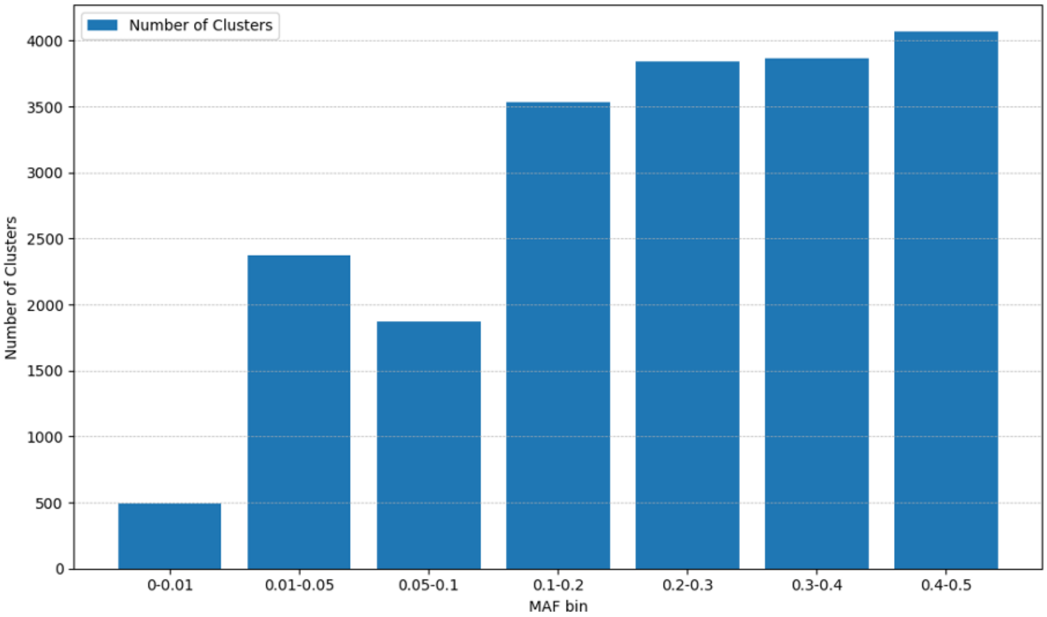

**Supplementary Figure 2.** Total number of colocalisation clusters by MAF bin. Error bars indicate 95% confidence intervals.

We quantified pleiotropy by (1) effect directionality, (2) absolute effect size distribution, and (3) the number of associated diseases per variant. Directional concordance — defined as the largest proportion of same-direction effects per variant (range = 0.5–1) — was 1 for non-pleiotropic SNPs. Among 9,828 pleiotropic variants, 1,793 (18%) showed concordance $<$ 1, indicating generally consistent effect directions.

The high directional concordance agrees with prior findings and reflects genetic correlations across diseases[11]. Nonetheless, 18% of variants showed discordant effects, likely due to noise, context-specific mechanisms, or discordant pleiotropy (opposing effects across diseases). The proportion of variants with discordant effects increased with MAF, which can be explained by several factors, including the prevalence of balanced or context-dependent effects in common variants, discordant pleiotropy, and statistical factors (Supplementary Figure 3).

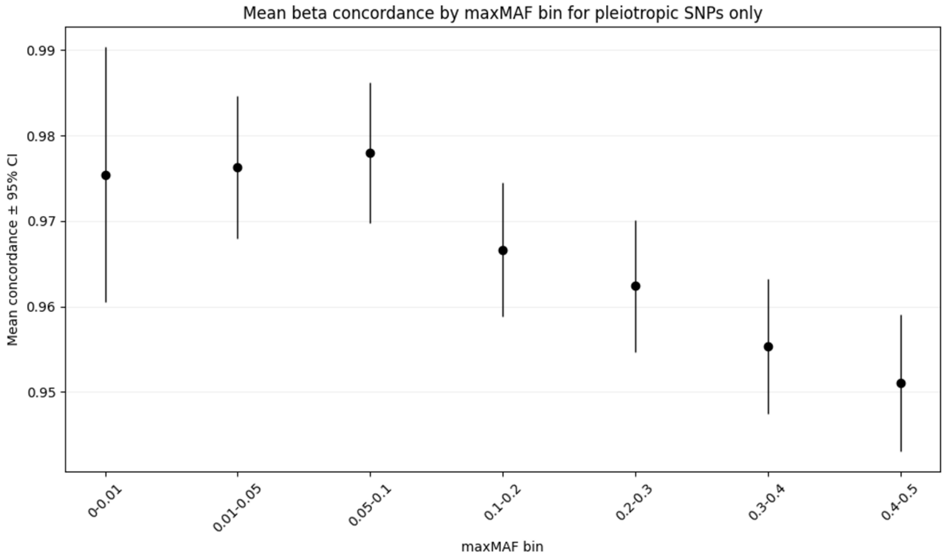

**Supplementary Figure 3.** Mean beta-effect concordance (proportion of same-direction associations across all trait pairs for a given variant) with 95% confidence intervals, plotted across MAF bins for pleiotropic lead variants only (vPS $>1$). Higher concordance indicates that the variant's effect on most associated traits points in the same direction; lower concordance indicates discordant pleiotropy.

We next examined effect size distributions among highly pleiotropic variants ($\geq$10 associated diseases). Because non-significant associations were set to zero, full distributions could not be assessed. For each of 118 variants, we compared single- and two-component Gaussian models of squared effect sizes using Akaike Information Criterion (AIC)/Bayesian Information Criterion (BIC); 100 (85%) favored the two-component model, indicating frequent bimodality (Supplementary Figure 4). On average, the smaller-effect mean was 14.5$\times$ lower than the larger-effect mean (median = 7.7), with $\sim$22% of associations belonging to the large-effect component. Thus, most associated traits exhibit modest effects, while a minority show much larger, likely core biological impacts. This bimodal pattern is consistent with the coexistence of direct biological effects on a minority of traits and smaller, indirectly mediated effects across the majority, likely reflecting the heterogeneous functional architecture of pleiotropic loci. This architecture is reminiscent of the omnigenic model[12], in which a small number of core genes exert direct effects on disease while a larger set of peripheral genes contributes smaller, trans-mediated effects.

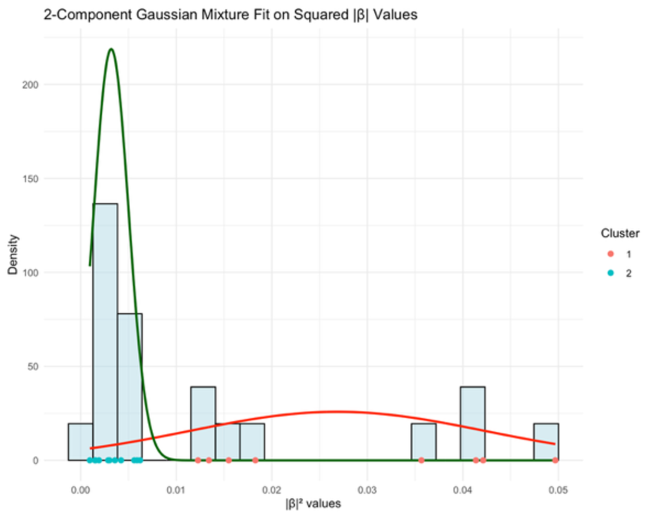

**Supplementary Figure 4.** Two-component Gaussian mixture model fitted to the distribution of squared absolute effect sizes ($|\hat{\beta}|^2$) across all lead variants. Cluster 1 (green) captures variants with small effect sizes; Cluster 2 (red) captures variants with large effect sizes. This bimodal decomposition was used to classify variants into large-effect and small-effect categories for downstream enrichment analyses.

Next, we analysed the number of traits associated per variant. Overall, 9,828 variants (24%) were pleiotropic (mean = 1.48; max = 85). The number of associations showed a non-linear relationship with MAF, increasing for both rare and common variants likely reflecting higher effect sizes in rare variants and greater detection power for common ones, consistent with prior findings. Given that most pleiotropic effects are small ($\approx$10–20$\times$ smaller than the maximal effect), we modeled the expected number of detectable associations assuming all traits are affected with effect sizes scaled by a factor of 11 relative to the maximum. Expected genome-wide significant associations ($P < 1 \times 10^{-8}$) were then estimated using each variant's maximal sample size, MAF, and corresponding non-centrality parameter (NCP).

The expected power to detect an association was computed as:

$$\text{Power} = P\left(\chi_1^2 > q_{\chi^2}(1 - 10^{-8}, df = 1) \mid ncp = \text{NCP}\right),$$

where

$$\text{NCP} = (\text{maxBeta}^2) \times \text{maxEffectiveSampleSize} \times 2 \times \text{maxMAF} \times (1 - \text{maxMAF})/11.$$

and where $q_{\chi^2}$ denotes the quantile of the chi-square distribution with one degree of freedom.

#### 7. gPS predicts trial safety independently of directional discordance

Gene-level beta-sign discordance was computed by aggregating variant-level beta-sign discordance, defined as 1$-$ concordance, across all L2G-prioritised variants per gene using both mean and maximum aggregation. Variants with no concordance information were assigned a concordance of 1 (discordance of 0). Three logistic regression models were fitted for trial safety gene membership: (1) $\log_2$(gPS) alone, (2) mean (or maximum) discordance alone, and (3) $\log_2$(gPS) and discordance jointly, each on $N$ = 8,285 disease-associated genes.

When using mean discordance, gPS retained its full predictive effect ($\beta$ = 0.29, $P$ = 5.4 $\times$ $10^{-11}$) while discordance was non-significant ($P$ = 0.83). Maximum discordance yielded the same pattern (gPS: $\beta$ = 0.25, $P = 5.9 \times 10^{-7}$; discordance: $P$ = 0.14), despite maximum discordance being significant in univariate analysis ($\beta$ = 1.82, $P = 7.8 \times 10^{-7}$), indicating that its predictive power is fully captured by gPS.

#### 8. Drug target enrichment bias due to sample size and therapeutic areas

We hypothesized that enrichment estimates could be biased or inflated due to uneven study representation across therapeutic areas (TAs), differences in sample sizes, or varying enrichment patterns between TAs. To address this, we extended the drug target enrichment framework to include multiple covariates. We implemented a mixed-effects logistic regression, modelling the effect of GWAS evidence and maximal study GWAS sample size for the disease as fixed effects and TA as a random effect. Analyses were performed on the full ChEMBL dataset to maximize statistical power. When the random TA effect was excluded, maximal sample size reduced the odds ratio (OR) of enrichment from 3.62 to 3.44, suggesting a small overestimation of enrichment. Including the random TA effect reduced the enrichment estimate to 3.14, and the variance of the TA component was estimated at 0.54 (SD = 0.74), indicating significant variability in enrichment across TAs. When only the random TA effect was included, excluding sample size, the OR decreased to 3.32, suggesting that TA and sample size had an independent impact on enrichment.

#### 9. Non-linearity of gPS vs drug target success

To thoroughly investigate the relationship between pleiotropy and the target outcome, we evaluated models incorporating linear and non-linear (quadratic) logarithmic pleiotropy terms. Pleiotropy was assessed using two metrics: the number of unique diseases (gPS) and the number of unique therapeutic areas (TAs).

For each metric, we compared three logistic regression models:

1. Model 0 (Baseline): Outcome $\sim$ Genetic Support
2. Model 1 (Linear): Outcome $\sim$ Genetic Support + log(X + 1)
3. Model 2 (Quadratic): Outcome $\sim$ Genetic Support + log(X + 1) + $\log(X + 1)^2$

##### 9.1. Likelihood Ratio Tests for Non-linearity

The inclusion of a quadratic logarithmic term (Model 2) significantly improved model fit over both the baseline and linear models, indicating a non-linear (inverted U-shaped) relationship.

- **Number of Therapeutic Areas (TAs):** The quadratic model vastly outperformed the baseline model (LR = 80.54, $P = 3.24 \times 10^{-18}$) and provided a significantly better fit than the linear model (LR = 64.90, df = 1, $P = 7.89 \times 10^{-16}$).
- **Number of Unique Diseases (gPS):** Similarly, the quadratic model outperformed the baseline model (LR = 66.38, $P = 3.85 \times 10^{-15}$) and the linear model (LR = 54.17, df = 1, $P = 1.84 \times 10^{-13}$).

##### 9.2. Robustness Checks via Bootstrapping

To ensure the quadratic effects were not driven by specific data compositions or outliers, we performed a stringent robustness check using 1,000 bootstrap iterations (resampling with replacement).

The negative quadratic term remained highly stable across all 1,000 iterations for both TAs and gPS metrics. Specifically, the quadratic coefficient maintained the exact same negative sign (100% sign stability) and remained statistically significant ($P < 0.05$) in 100% of the iterations.

##### 9.3. Bootstrap Robustness Metrics (1,000 Iterations)

For models assessing the number of Therapeutic Areas (TAs), the original linear logarithmic coefficient was 0.568, with a bootstrapped mean of 0.568 (Std: 0.064) and a 95% confidence interval (CI) of [0.441, 0.692]. The original quadratic term was -0.267, with a bootstrapped mean of -0.266 (Std: 0.034) and a 95% CI of [-0.332, -0.198]. Both terms were significant ($P < 0.05$) in 100% of the bootstrap iterations.

Similarly, for models assessing the number of Unique Diseases (gPS), the original linear logarithmic coefficient was 0.371, with a bootstrapped mean of 0.370 (Std: 0.047) and a 95% CI of [0.271, 0.459]. The original quadratic term was -0.120, with a bootstrapped mean of -0.120 (Std: 0.017) and a 95% CI of [-0.152, -0.085]. Again, both terms achieved a 100% significance rate across all iterations.

##### 9.4. Sensitivity Analysis: Exclusion of Safety Liabilities

To ensure the observed non-linear effect was not driven by targets with known safety issues, we performed an additional sensitivity analysis by excluding all targets associated with documented safety liabilities (specifically: 'trial_safety_concern', 'withdrawn_drug', and 'liable_target'). Filtering these targets reduced the dataset from 37,377 to 8,433 observations. Despite the reduced sample size, the quadratic relationship remained robust and highly significant. For the therapeutic areas (TAs) metric, the quadratic logarithmic term retained a strongly negative and highly significant coefficient (-0.474, $P < 0.001$), demonstrating that the inverted U-shaped pattern is robust and not an artifact of safety-related attrition.

#### 10. Phase transition probabilities stratified by pleiotropy

To examine whether gene pleiotropy influences the probability of advancing through clinical development phases, we analysed phase-to-phase transition rates for all 37,377 target–indication (T–I) pairs in the ChEMBL dataset, stratified by pleiotropy level. Pleiotropy was operationalised as the number of unique therapeutic areas (TAs) associated with the target gene and binned into three groups: Low (1 TA), Medium (2–5 TAs), and High ($\geq$ 6 TAs). T–I pairs with no GWAS genetic support (uniqueTherapeuticAreas = 0) were excluded from this stratified analysis, leaving 18,480 T–I pairs (Low: $n$ = 6,578; Medium: $n$ = 9,705; High: $n$ = 2,197).

For each pleiotropy group and each of the three clinical transitions (Phase I$\to$II, Phase II$\to$III, Phase III$\to$ approval), the transition probability was estimated as the proportion of T–I pairs that reached at least the end phase among those that reached at least the start phase:

$$P(\text{transition}) = \frac{|\{i : \text{maxPhase}_i \geq \text{phase}_{\text{end}}\}|}{|\{i : \text{maxPhase}_i \geq \text{phase}_{\text{start}}\}|}.$$

This cross-sectional attrition approach includes currently active pairs at each phase alongside completed ones, providing a snapshot estimate consistent with prior analyses of clinical success rates[13]. Wilson 95% confidence intervals were computed for each proportion.

Statistical significance was assessed using a chi-square omnibus test for each transition, followed by pairwise two-sided proportions $z$-tests (Medium vs Low, High vs Low, High vs Medium). $P$-values from all nine pairwise tests were adjusted jointly using the Benjamini–Hochberg (BH) false discovery rate procedure. Relative risks (RRs) with 95% confidence intervals were computed via the delta method applied to the log-RR:

$$\widehat{RR} = \frac{\hat{p}_a}{\hat{p}_b}, \qquad \mathrm{SE}(\log \widehat{RR}) = \sqrt{\frac{1-\hat{p}_a}{n_a\hat{p}_a} + \frac{1-\hat{p}_b}{n_b\hat{p}_b}}.$$

Results are summarised in Supplementary Materials Table 2. Phase II→III was the only transition showing significant differences across pleiotropy groups (omnibus $\chi^2$, $P < 0.001$). Transition rates at Phase I→II (82.6–84.2%) and Phase III→approval (29.4–31.2%) did not differ significantly after FDR correction. At Phase II→III, Low-pleiotropy targets showed the highest transition rate (57.0%), followed by Medium (53.5%) and High (49.8%). All three pairwise comparisons at this transition remained significant after BH correction: Medium vs Low (RR = 0.94, 95% CI 0.91–0.97, $P_{\mathrm{adj}} = 3\times10^{-4}$), High vs Low (RR = 0.87, 95% CI 0.83–0.92, $P_{\mathrm{adj}} < 0.001$), and High vs Medium (RR = 0.93, 95% CI 0.88–0.98, $P_{\mathrm{adj}} = 0.010$).

**Supplementary Materials Table 2.** Phase transition success rates and pairwise relative risks by pleiotropy level. $N$ = number of T–I pairs entering each transition. Success % = proportion advancing to the next phase. RRs and 95% CIs are computed via the log-RR delta method. $P_{\mathrm{adj}}$ = BH-FDR corrected $P$-value across all nine pairwise tests. Significant comparisons ($P_{\mathrm{adj}} < 0.05$) are shown in bold.

| Transition | Pleiotropy | $N$ | Success % | Omnibus $P$ | Comparison | RR (95% CI) | $P_{\mathrm{adj}}$ |
|---|---|---|---|---|---|---|---|
| Phase I→II | Low (1 TA) | 6,578 | 82.9% | 0.038 | Med vs Low | 1.02 (1.00–1.03) | 0.058 |
| | Medium (2–5) | 9,705 | 84.2% | | High vs Low | 1.00 (0.97–1.02) | 0.759 |
| | High (≥6) | 2,197 | 82.6% | | High vs Med | 0.98 (0.96–1.00) | 0.117 |
| Phase II→III | Low (1 TA) | 5,453 | 57.0% | <0.001 | **Med vs Low** | **0.94 (0.91–0.97)** | $3\times10^{-4}$ |
| | Medium (2–5) | 8,173 | 53.5% | | **High vs Low** | **0.87 (0.83–0.92)** | <**0.001** |
| | High (≥6) | 1,815 | 49.8% | | **High vs Med** | **0.93 (0.88–0.98)** | **0.010** |
| Phase III→App. | Low (1 TA) | 3,110 | 29.4% | 0.238 | Med vs Low | 1.06 (0.99–1.14) | 0.136 |
| | Medium (2–5) | 4,376 | 31.2% | | High vs Low | 1.04 (0.93–1.16) | 0.638 |
| | High (≥6) | 903 | 30.6% | | High vs Med | 0.98 (0.88–1.09) | 0.759 |

#### 11. Integration to Open Targets Platform

The Gentropy framework was designed to be fully reproducible, allowing this effort to be ongoing rather than a one-off analysis. All generated data and pipelines have been integrated into the Open Targets ecosystem and are accessible through the Open Targets Platform[14]. We introduced three new interfaces that capture complementary dimensions of the dataset: a variant page summarising molecular and complex-trait associations, external annotations (ClinVar/ClinGen, UniProt, Pharmacogenetics); a credible-set page displaying fine-mapping results, colocalisations and L2G predictions; and a study page providing harmonised study-level metadata and associations. L2G scores are accompanied by SHAP-based explanations to facilitate interpretability (Supplementary Methods). GWAS-derived evidence is aggregated into direct disease-level associations using harmonic sum and further propagated using the ontology to generate indirect associations (Supplementary Methods). The Open Targets Platform and data are updated quarterly, with ongoing expansion of GWAS coverage and continuous refinement of the target-prioritisation framework, ensuring results remain current with the latest available GWAS evidence.

All data and large-scale fine-mapping results are released under a CC0 licence to facilitate drug target discovery and research in statistical genetics of complex traits. The Open Targets Platform provides open access to these data sets (`https://platform.opentargets.org/`). The scientific community continues to provide extensive benchmarking and evaluation of the resource.

### Supplementary Methods

Below we describe the Gentropy pipeline overview and orchestration, data sources, and statistical methods used to obtain the complex statistical genetics evidence that was provided to the user interface of the Open Targets Platform (indicated by the datasource: GWAS associations).

#### 1. Orchestration

The Gentropy pipelines consist of several parts, detailed in Sections 3, 4 and 5 below and depicted in Supplementary Figure 5.

The first part was preparation of static assets that were used further for the analysis. This part prepared GnomAD variant annotation and linkage disequilibrium (LD) matrices/indices to be further used by the fine-mapping pipelines.

Next, a datasource-specific Gentropy pipeline part ingested the input raw summary statistics, curated SNP–trait associations or results of fine-mapping from external data sources and transformed them into Gentropy datasets. Each datasource-specific Gentropy pipeline generated two datasets: `StudyIndex` and `StudyLocus`.

The last part of the pipelines was the Genetics ETL that is also part of the Open Targets unified pipeline. It used the previously generated **StudyIndex** and **StudyLocus** datasets as input. Within the Genetics ETL, we performed the validation of studies and credible sets, generated the variant index, ran colocalisation, generated the L2G feature matrix and ran the L2G prediction.

The reason to separate the Genetics ETL from the datasource-specific Gentropy pipeline is to avoid recalculation of the computationally expensive steps of fine-mapping.

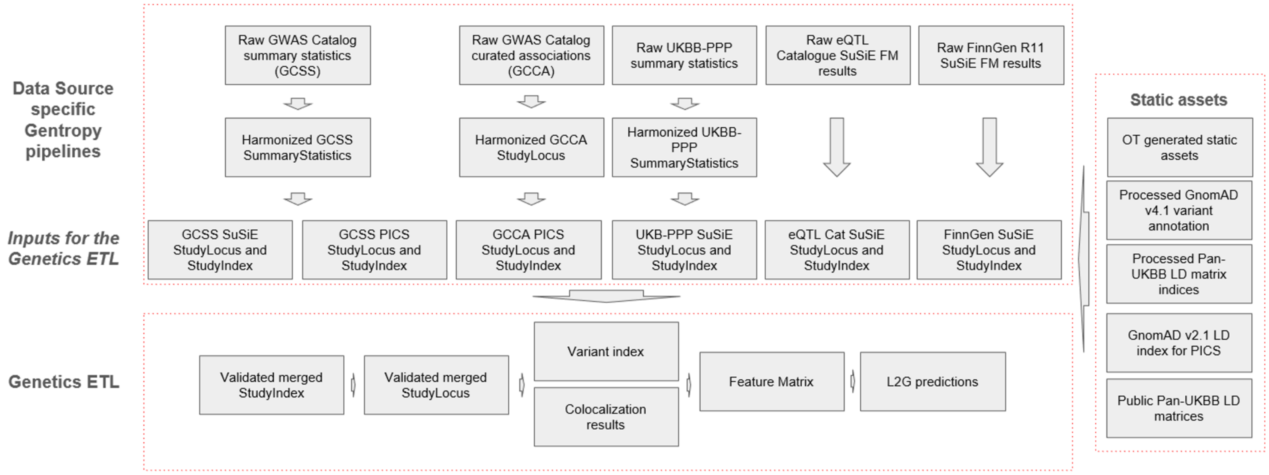

**Supplementary Figure 5.** Schematic overview of the Gentropy data processing pipelines. The top layer shows data source-specific pipelines harmonising raw input data from five sources (GWAS Catalog summary statistics, GWAS Catalog curated associations, UKB-PPP, eQTL Catalogue, and FinnGen) into standardised StudyLocus and StudyIndex objects. The middle layer shows the resulting inputs to the Genetics ETL alongside static reference assets. The bottom layer shows the Genetics ETL outputs: a validated merged StudyIndex, merged StudyLocus, variant index, colocalisation results, feature matrix, and L2G predictions.

#### 2. Data sources

We used different data sources throughout the pipeline. Each data source is described in its own section. The data sources are divided into two groups: (1) *static assets*—datasets used for fine-mapping and for different annotations, these assets are not usually expected to change much from release to release; and (2) *association datasets*, described in Section 2.2.

##### 2.1. Static assets

**GnomAD variant annotation and LD.** GnomAD (Genome Aggregation Database)[15] is a comprehensive resource that provides aggregated genomic data from large-scale sequencing projects. It encompasses variants from diverse populations and is widely used for variant annotation and population genetics studies.

We processed the GnomAD v4.1 variant annotation file hail table from the GnomAD public Google Cloud bucket. It was used as the reference for UKB-PPP GWAS summary statistics unification and for the variant index in Genetics ETL.

We used GnomAD v2.1.1 LD matrices[16] to create the LD index that is used for PICS fine-mapping (see below). In short, the LD index is the list of all variants presented in the LD matrix with its corresponding proxies with $r^2 \geq 0.5$ for all GnomAD v2.1.1 populations; however, only five populations—African-American (AFR), American Admixed/Latino (AMR), East Asian (EAS), Finnish (FIN), and Non-Finnish European (NFE)—were used for PICS fine-mapping.

**Pan-UKBB LD matrices.** We used Pan-UK Biobank project[17] LD matrices for three ancestries: Non-Finnish European (NFE), Central/South Asian (CSA) and African (AFR). The LD was computed for each chromosome in a 10 Mb radius. Only SNPs with INFO $> 0.8$ and MAC $> 20$ in each population were used to calculate LD. Matrices are stored in Hail BlockMatrix format. We used these LD matrices for SuSiE fine-mapping.

**Other static assets.** We used the following objects generated by the Open Targets Platform ETL: target index, disease index, biosample index, and protein–protein interactions (StringDB, https://string-db.org/).

##### 2.2. Association datasource specific Gentropy pipelines

**Overview.** The result of each datasource-specific pipeline is `StudyIndex` and `StudyLocus`. Depending on the datasource the pipelines can also generate the harmonised `SummaryStatistics` object as an intermediate output. Where applicable, we performed datasource-specific fine-mapping. For each data source we applied specific filtering and/or flag assignment that was further used for validation in Genetics ETL.

**The NHGRI-EBI GWAS Catalog.** The NHGRI-EBI GWAS Catalog[4] is a datasource maintained at EMBL-EBI that provides detailed, structured genome-wide association study data in standardised format of summary statistics and curated associations (top-hits) to EFO traits, providing a rich source of genetic associations. We used GWAS Catalog to produce three sets of `StudyLocus` and `StudyIndex` datasets:

1. **GWAS Catalog curated associations** PICS fine-mapping results (**GCCA** PICS).
2. **GWAS Catalog summary statistics** PICS fine-mapping results (**GCSS** PICS).
3. **GWAS Catalog summary statistics** SuSiE fine-mapping results (**GCSS** SuSiE).

*GCSS unification and harmonisation.* For each study, we performed additional harmonisation and converted it to the Gentropy `SummaryStatistics` format. Additional harmonisation and quality control included: (1) filtering out SNPs with unavailable beta, standard error or *p*-value; (2) filtering out SNPs with zero or Inf values for beta and standard error; (3) filtering out SNPs with negative *p*-value or standard error; (4) filtering out SNPs with *p*-value equal to 1. In some cases, this resulted in empty GWAS files and these studies were excluded from further analysis. If available, ancestries from the GWAS Catalog were mapped to GnomAD ancestry suffixes. The ancestry was not assigned if it was unavailable or not present in the dictionary.

*GCSS studies manual curation.* We manually curated the GCSS studies from the file "All studies – With study accession numbers, ontology annotations, genotyping technology, cohort identifiers and full summary statistics availability" which had a corresponding summary statistics file in the GWAS Catalog FTP site (even if it was empty). Based on the corresponding publication we assessed the following:

1. Specific study type—whether pQTL, microbiome GWAS or all other trait GWAS. Studies identified as pQTL and microbiome GWAS were excluded from further analysis.
2. Specific analysis flag—where possible we assigned the following analysis flags to the study: multivariate analysis, ExWAS, non-additive model, metabolite, GxG, GxE, case-case study. If the study was flagged by any of the flags except "metabolite" it was excluded from SuSiE fine-mapping and was fine-mapped by PICS due to incompatibility with the SuSiE fine-mapping analysis type.

*GCCA unification and harmonisation.* We harmonised GWAS Catalog curated associations from the table "All associations – with added ontology annotations, GWAS Catalog study accession numbers and genotyping technology". The GWAS Catalog curation process may have resulted in multiple GWAS being grouped under one study accession, in which case one study accession is split into multiple studyIDs. For example, if the reported trait is 'Obesity-related traits' but the underlying association data informs which top-hit comes from which specific trait, we created as many new studies as there were association-level trait annotations with the new study identifiers generated from the original study accession plus a suffix. A similar action is required when results from different ancestries are pooled under one study accession.

The harmonisation process included a number of steps that flag any associations with quality concerns. When harmonising GCCA, the following cases were flagged:

- Variant interaction associations.
- Variant location is not available in an association source.
- Variant location data is inconsistent (chromosome and position values do not match).
- Lead variant is duplicated for the same study.
- The provided variant location does not match any GnomAD v4.1 variant.
- The risk allele could not be mapped to the GnomAD v4.1 reference.
- The lead variant had palindromic alleles.
- The lead variant *p*-value was above the genome-wide significant threshold ($p > 10^{-8}$).

Curated associations were converted into the Gentropy `StudyLocus` format and uniformly flagged with "Study locus from curated top hit".

*Clumping and fine-mapping of GCSS and GCCA PICS.* We applied distance and LD-based clumping to all available GCSS and GCCA. Details of the methods are given in Section 4. In short, we used a *p*-value threshold of $\leq 10^{-8}$ and 500,000 bp as a radius for distance-based clumping. We applied PICS fine-mapping[18,1] to the clumping results using the LD index based on GnomAD v2.1.1. In the case of multiple-ancestry studies, we used Non-Finnish Europeans (NFE) if it was in the list of ancestries, or major ancestry otherwise. If the lead variant was not present in the LD index, we assumed it was a credible set with the size of one (credible set with only lead variant) but flagged this `StudyLocus` with “Variant not found in LD reference”. We did not produce fine-mapping results if the study’s ancestry was not in the list of available LD index ancestries. All resulting `StudyLocus` objects were additionally flagged with “Study locus fine-mapped without in-sample LD reference”.

*Clumping and fine-mapping of GCSS SuSiE.* To form the `StudyLoci` for fine-mapping, we applied the locus breaker method (see Section 4.2) to all available GCSS using the default parameters and lead SNP $P \leq 10^{-8}$. We performed SuSiE fine-mapping[2] only on StudyLoci that met the following criteria:

1. Major ancestry for the study is NFE, AFR, CSA or EAS. If the major ancestry was EAS, we used CSA instead.
2. The study type is “gwas”.
3. The study has no analysis flags except “metabolite”.
4. Study has no quality control flags (as described above).
5. The locus did not overlap with the MHC region. We did not exclude X or Y chromosomes if they were present in GWAS.
6. The number of SNPs in the locus after overlapping with the LD matrix was in the range [100, 15,000].

For all eligible study loci, we applied the SuSiE-inf method without estimation of infinitesimal effects (equivalent to the classical SuSiE method) using Pan-UKBB LD matrices. We performed filtering of the resulting 95% CSs based on CS log(BF) $\leq 2$, minimum $r^2$ purity $\leq 0.25$ and lead SNP $P \geq 10^{-5}$. Additionally, within a locus we checked the pairwise $r^2$ between lead variants and removed one of the CSs if $r^2 \geq 0.8$, retaining the CS with the more significant $P$. Where the locus breaker procedure created overlapping loci, we could obtain credible sets with duplicated lead variants within the same study; if two CSs from different loci within a study had the same lead variant, we removed one, retaining the CS with the largest CS log(BF). All resulting `StudyLocus` objects were additionally flagged with “Study locus fine-mapped without in-sample LD reference”.

**eQTL Catalogue.** The eQTL Catalogue[6] provides unified gene expression, protein level and splicing QTLs from available public human studies. It serves as the key resource of molecular QTLs (molQTLs) that we used for colocalisation and target prioritisation. The eQTL Catalogue uses the QTLmap pipeline[6] for molQTL mapping with SuSiE as the fine-mapping method, resulting in 95% credible sets (CSs).

We transferred the raw fine-mapping results from the eQTL Catalogue FTP server to the OT Google buckets and converted all available CSs and study information to Gentropy `StudyIndex` and `StudyLocus` objects. We flagged CSs with lead SNP $P \geq 10^{-3}$ with the “Subsignificant $P$” flag.

**FinnGen.** FinnGen[5] is an academia-industry partnership that aims to produce genome variant data for 500,000 Finns. The genomic data is combined with phenotype data collected by national health registries.

Because the Finnish population has been genetically isolated, fine-mapping is performed with a suitable reference panel of LD. The FinnGen study has performed fine-mapping using SuSiE and FINEMAP, based on a reference panel of whole genome sequencing data from Finns.

We used the latest **FinnGen R12**. We accessed the study information and results of the SuSiE fine-mapping for 95% CSs and converted them to Gentropy `StudyIndex` and `StudyLocus` objects. We flagged CSs with lead SNP $P \geq 10^{-5}$ with the "Subsignificant *P*" flag.

**The UK Biobank Pharma Proteomics Project (UKB-PPP).** The Pharma Proteomics Project is a pre-competitive biopharmaceutical consortium characterising the plasma proteomic profiles of 54,219 participants in the UK Biobank[7].

*GWAS harmonisation.* We used GWAS results for 2,954 proteins of European ancestry. The original GWAS were downloaded from the Synapse platform. The GWAS were further unified in Gentropy `SummaryStatistics` format. SNPs with MAF $< 10^{-4}$ and INFO $< 0.8$ were filtered out. We also unified the order of effective and reference alleles with the GnomAD annotation—if the alleles were reversed, we changed the sign of the effect size; if the allele combination did not match the reference, we filtered out the SNP.

*SuSiE fine-mapping.* To form the `StudyLoci` for fine-mapping, we applied the locus breaker method to all available UKB-PPP studies using the default parameters and lead SNP $P \leq 1.7 \times 10^{-11}$. For all resulting `StudyLocus` objects, we applied the SuSiE-inf method without estimation of infinitesimal effects using Pan-UKBB LD matrices for the EUR population. We performed filtering of the resulting 95% CSs in a similar way to the GCSS SuSiE procedure described above. All resulting `StudyLocus` objects were additionally flagged with "Study locus fine-mapped without in-sample LD reference".

#### 3. Genetics ETL

`StudyLocus` and `StudyIndex` objects from each of the six datasource-specific Gentropy pipelines described above were sourced as input to the Genetics ETL.

##### 3.1. Study validation

The aim of the study validation step is to unify and validate the individual `StudyIndex` datasets generated by Gentropy datasource-specific pipelines. The step returns a single `StudyIndex` dataset that consists of valid and unique studies. The following data sources are used for study validation: (1) GCCA study index, (2) GCSS PICS study index, (3) GCSS SuSiE study index, (4) eQTL Catalogue study index, (5) UKB-PPP study index, and (6) FinnGen study index.

To avoid duplication in the `StudyIndex` records coming from GWAS Catalog, we carried out the **deconvolution of studies based on the availability of summary statistics and quality control flags**. The study validation is based on a flagging system. We joined all indexes together and applied the following procedure of flag assignment:

1. If `studyId` was duplicated:
   (a) Concatenate all `studyTypes` if different within duplicated rows.
   (b) Concatenate all available flags in `qualityControls` and `analysisFlags` within duplicate rows.
   (c) Flag all duplicated rows except the first.

2. Flag studies with non-supported study types (`gwas`, `(sc)eqtl`, `pqtl`, `(sc)sqtl`, `(sc)tuqtl`).
3. Flag molQTL studies with invalid `targetId`.
4. Flag complex trait studies with invalid trait ontologies.
5. Flag molQTL studies with invalid biosample.

We removed all studies that had at least one of the following flags:

1. Target/gene identifier could not match to reference.
2. No valid disease identifier found.
3. This type of study is not supported.
4. Biosample identifier was not found in the reference.
5. The identifier of this study is not unique.
6. The mean beta QC check value is not within the expected range.
7. The PZ QC check values are not within the expected range.
8. The GC $\lambda$ value is not within the expected range.
9. GWAS Catalog study has not been curated by Open Targets (except when the flag "Harmonised summary statistics are not available or empty" was present at the same time).

##### 3.2. StudyLocus validation

Study locus validation (credible sets) is also based on a flagging system. We merged all `StudyLocus` objects from the same data sources as for study validation. We filtered the SNPs in `StudyLocus` to the minimum number of SNPs to form a 95% CS. We assigned the appropriate flag to the CS if it had one of the following characteristics:

1. The lead variant is within the MHC region (chr6:25,726,063–33,400,556).
2. The lead variant has a chromosome identifier outside the valid set (1–22, X, Y, XY, mt).
3. The CS study is not in the list of valid studies.
4. It is the GCSS PICS CS and it has a valid SuSiE CS from the same region and study.
5. It is the GCCA PICS CS and it has a valid PICS CS from the same region and study.
6. The sum of PIPs in the CS was not within [0.95, 1].

We removed all `StudyLocus` objects that had at least one of the flags listed in the pipeline configuration, ensuring that the resultant merged `StudyLocus` object consists of only unique and valid credible sets.

##### 3.3. Variant index and variant annotation

To build a comprehensive representation of variants in the Open Targets Platform, we compiled a list of variants—the "variant index". It was defined as any variant for which phenotypic information was available. The index contained all unique variants from all valid credible sets, curated disease-related variants from ClinVar and UniProt, and variants for which pharmacogenomics information is available from PharmGKB. Note that downstream pleiotropy analyses used only variants from valid credible sets. Variants were then annotated via Ensembl's Variant Effect Predictor (VEP)[19,20].

Annotated variant index features include:

1. Transcript consequences are required in downstream locus-to-gene prediction. For this purpose, we collected distances from footprint and transcription start site (TSS) only for Ensembl canonical transcripts with a distance $>$500 kbp. Annotations from a number of open source *in silico* prediction methods were harvested to inform conservation (GERP)[21], potential loss of function (LOFTEE)[16], and impact on protein folding (AlphaMissense)[22].
2. Overlapping variants provided by VEP helped us enrich annotation data with rs identifiers and cross-references to ClinVar.

Allele frequencies were extracted from GnomAD v4.1 joint genome and exome datasets directly[15]. The FoldX dataset[23] provides free energy changes of the amino acid substitution caused by missense variants.

A short description of the applied methods to assess variant effect is given in Supplementary Materials Table 3.

**Supplementary Materials Table 3.** Summary of variant effect prediction methods used for variant annotation.

| Method | Description |
|---|---|
| AlphaMissense[22] | A deep learning model that builds on the protein structure prediction tool AlphaFold2 to assess the effect of missense variants across the proteome. |
| FoldX[23] | A computational tool that predicts the impact of mutations on protein stability and structure by calculating changes in free energy, helping to assess the potential functional consequences of variants. |
| GERP[21] | GERP (Genomic Evolutionary Rate Profiling) scores are used to identify regions of the genome that are evolutionarily conserved and likely to be functionally important, with higher scores indicating potential deleterious impact of variants. |
| LOFTEE[16] | LOFTEE (Loss-Of-Function Transcript Effect Estimator) identifies and annotates high-confidence loss-of-function variants in human genetic data, focusing on variants that likely disrupt gene function. |
| SIFT[24] | SIFT (Sorting Intolerant From Tolerant) predicts whether an amino acid substitution affects protein function based on sequence homology and the physical properties of amino acids. |
| VEP[19,20] | Pathogenicity score derived from the most severe consequence term provided by Ensembl's Variant Effect Predictor (VEP). |

##### 3.4. Calculation of StudyLocus overlaps and colocalisation

Prior to colocalisation, we calculated the overlaps between `StudyLocus` objects. The overlap is defined as a pair of 95% credible sets with at least one SNP in common. We calculated overlaps between all GWAS study loci and between GWAS and molQTL study loci. We did not calculate overlaps between molQTL CSs. For all resulting overlaps we applied colocalisation and colocalisation directionality assessment (see Section 4.7 and 4.8).

##### 3.5. Feature matrix generation

Validated credible sets, the results of colocalisation, variant index and gene index were used to generate a feature matrix—the table of functional genomics features for each of the genes around the credible sets. The description of features is available in the main Methods section. The feature matrix was saved in a wide format, having dimensions [Number of all genes around all CSs $\times$ Number of features]. The subset of rows from the feature matrix was used to train the L2G model. We generated the feature matrix for both protein-coding and non-protein-coding genes.

##### 3.6. L2G predictions

The "locus-to-gene" (L2G) machine learning model prioritises likely causal genes at each GWAS locus by using the functional genomics features. We applied a pre-trained L2G model (see main Methods section) to all rows in the feature matrix. Resulting L2G scores were filtered to protein-coding genes only and by score $\geq$ 0.05. It was used to calculate Open Targets Platform evidence, direct and indirect associations.

##### 3.7. L2G explanations

Using the SHAP (SHapley Additive exPlanations) library, we extracted feature importance values for all L2G predictions—Shapley values. Shapley values provide a principled approach based on game theory to explain the contribution of individual features or groups of features, revealing how each group influences the final L2G score. These contributions are approximated to be additive, meaning the sum of the Shapley values for all feature groups equals the total L2G score.

We aggregated the Shapley values into the following meaningful feature groups:

- *Base Shapley value*: represents the baseline before any feature-specific information is considered, and is therefore equivalent for all genes and credible sets.
- *Distance-based features*.
- *Colocalisation-based features*.
- *VEP (Variant Effect Predictor)-based features*.

This approach helps identify which types of evidence (e.g., distance, colocalisation, or functional impact) are most influential for a given locus-to-gene association.

#### 4. Statistical methods

##### 4.1. GWAS summary statistics quality control

We performed GWAS summary statistics quality control (QC) for GWAS Catalog studies with available summary statistics using established methods[25]. We applied the following methods:

1. **The P–Z test.** We checked the difference between the log *p*-values reported in the study and those derived from the reported betas and standard errors. We checked the mean and standard deviation of the difference. If at least one value for the study was greater than 0.05, we considered the study to have failed QC and flagged it with "The PZ QC check values are not within the expected range".
2. **The mean beta check.** We checked the mean value of the beta across all SNPs in the study. If the absolute mean value exceeded 0.05 we considered the study to have failed QC and flagged it with "The mean beta QC check value is not within the expected range".
3. **The genomic control (GC) $\lambda$ check.** We calculated the additive GC $\lambda$ for all studies. If the GC $\lambda$ value was not within the [0.7, 2.5] range we considered the study to have failed QC and flagged it with "The GC $\lambda$ value is not within the expected range".

Additionally, we flagged all GWAS studies that have fewer than 2,000,000 SNPs with "The number of SNPs in the study is below the expected threshold". This flag does not indicate that a study failed QC but is used to indicate that the study is not eligible for SuSiE fine-mapping.

##### 4.2. Clumping

Clumping is a technique for selecting the most significant SNP within a region or set of SNPs in LD, essentially removing redundant signals and focusing on the most likely causal variant in that region. We used three methods of clumping in our pipelines.

**Distance-based clumping.** The method is based on an iterative procedure in which the variant with the strongest *p*-value is selected and all other variants within a predefined distance (radius) from this SNP are clumped together. The procedure is repeated as long as there is at least one significant SNP. The output of the method is a list of lead variants. The two parameters are the distance to clump and the *p*-value significance threshold.

**LD-based clumping.** The method is applied to the results of the distance-based clumping (list of lead variants). It is again based on an iterative procedure in which the variant with the strongest *p*-value is selected and all other variants with high LD ($r^2 \geq 0.5$) are clumped together. We used the pre-calculated LD index (see above) to infer the variants in high LD with the lead variant. We used both distance- and LD-based clumping to define loci for PICS fine-mapping of GCSS and GCCA.

**Locus breaker.** The method consists of three steps:

1. In the first step, we performed regular distance-based clumping.
2. In the second step, we filtered the input summary statistics by the baseline *p*-value (default $10^{-5}$). We then clumped SNPs that are closer to each other than the cut-off distance (default 250,000 bp). Next, we filtered clumps by having at least one variant with a *p*-value above the genome-wide significance threshold. At this stage, we define a list of loci consisting of information about the lead variant and the locus boundaries (the most left and right SNPs in the clump). To each of the locus boundaries we subtract/add the flanking distance (default 100,000 bp) to avoid situations where the locus size consisting of only one lead variant is 0.

3. In the third step we selected the loci with a size greater than the specified large locus size threshold (1,500,000 bp by default) and “break” each of the large loci using the lead variants from the distance-based clump that lie within the boundaries of the large locus. For each of the distance-based lead variants, we assign the boundaries as $\pm$half the size of the large locus size threshold. Thus, each large locus is divided by several overlapping loci of the size of the large locus size threshold. The small loci are unaffected by the splitting.

The general procedure results in a list that contains information about the lead variant and the locus boundaries (the most left and right SNPs in the cluster), and the largest locus size does not exceed the large locus size threshold. The boundaries of the locus are used to define the region and are used in SuSiE fine-mapping for LD matrix ingestion. The locus breaker results in much smaller locus sizes on average compared to those defined by window-based clumping.

##### 4.3. PICS fine-mapping

The PICS algorithm was originally implemented[18] for investigating the fine-mapping of causal autoimmune disease variants. It is a method to fine-map the most likely causal SNPs associated with a trait or disease within a haplotype. The algorithm is based on the calculation of the Posterior Inclusion Probabilities (PIP) of tag variants linked to the lead variant by LD within the target population. We reimplemented[1] the original algorithm using PySpark. The calculation is performed using all proxy variants with $r^2 > 0.5$ and the default parameter $k = 6.4$ as reported in the original paper. Our implementation uses a precomputed LD index (see above) to define the list of SNPs in high LD with the lead variant.

##### 4.4. SuSiE fine-mapping

If both summary statistics and high-precision LD are available for the locus, we used SuSiE-inf fine-mapping[2]. This method is a generalisation of the original SuSiE, allowing modelling of infinitesimal effects alongside fewer larger causal effects.

SuSiE-inf has two approaches for updating estimates of the variance components—Method of Moments and Maximum Likelihood Estimator (MoM/MLE). The function takes an array of $Z$-scores and a matrix of variant LD to perform fine-mapping. There is a boolean option `est_tausq` that enables the estimation of infinitesimal effects; if it is disabled, it performs fine-mapping equivalent to the original SuSiE method. We used SuSiE-inf only in combination with Pan-UK Biobank LD matrices for three ancestries: NFE, CSA and AFR.

##### 4.5. CARMA outlier detection

CARMA is a method for fine-mapping and outlier detection. We implemented a simplified version of CARMA with the following features: (1) it uses only Spike-slab effect size priors and Poisson model priors; (2) the C++ core is re-implemented in Python; (3) the way of storing the configuration list is changed, using a string with the list of indices for causal SNPs instead of a sparse matrix; (4) fixed bugs in PIP calculation; (5) no credible models or credible sets, only PIPs; (6) no functional annotations; (7) removed unnecessary parameters. We did not use CARMA for outlier detection in the 25.06 release.

##### 4.6. Summary statistics imputation

Summary statistics imputation leverages LD information to compute $Z$-scores of missing SNPs from neighbouring observed SNPs by taking advantage of LD. We implemented the basic model from the RAISS (Robust and

Accurate Imputation from Summary Statistics) package[26]. We did not use summary statistics imputation in the 25.06 release.

##### 4.7. COLOC colocalisation

If both CSs in the overlap had log(BF) information (both CSs are results of SuSiE fine-mapping) we conducted colocalisation analysis using the COLOC method[27]. COLOC is a Bayesian method which, for two traits, integrates evidence over all variants at a locus to evaluate the following hypotheses:

- $H_0$: No association with either trait.
- $H_1$: Association with trait 1, not with trait 2.
- $H_2$: Association with trait 2, not with trait 1.
- $H_3$: Association with trait 1 and trait 2, two independent SNPs.
- $H_4$: Association with trait 1 and trait 2, one shared SNP.

The process to compute the posterior probabilities of the hypotheses is as follows. Given Bayes Factor (BF) columns with null values filled to 0 and the log Bayes Factors (logBF) summed for overlapping signals, we aggregate BF vectors for each pair of signals by study locus:

- Compute $\text{logsum}_1$: log-sum of BFs for trait 1.
- Compute $\text{logsum}_2$: log-sum of BFs for trait 2.
- Compute $\text{logsum}_{12}$: log-sum of combined BFs for traits 1 and 2.

Adding priors and calculating BFs for hypotheses:

- $H_0$ (no association): $\log \text{BF} = 0$.
- $H_1$ (trait 1 only): $\log \text{BF} = \log(p_{c1}) + \text{logsum}_1$.
- $H_2$ (trait 2 only): $\log \text{BF} = \log(p_{c2}) + \text{logsum}_2$.
- $H_3$ (both traits, different causal variants):

$$\begin{aligned}
\text{sumlogsum} &= \text{logsum}_1 + \text{logsum}_2 \\
M &= \max(\text{sumlogsum}, \text{logsum}_{12}) \\
\log \text{diff} &= M + \log(\exp(\text{logsum}_1 - M) - \exp(\text{logsum}_{12} - M)) \\
\log \text{BF} &= \log(p_{c1}) + \log(p_{c2}) + \log \text{diff}
\end{aligned}$$

- $H_4$ (both traits, shared causal variant): $\log \text{BF} = \log(p_{c12}) + \text{logsum}_{12}$.

Posterior probabilities for $H_0$–$H_4$ are obtained by normalising the combined logBF vector: $P(H_i) = \exp(\log \text{BF}_i - \log \sum_j \exp(\log \text{BF}_j))$.

##### 4.8. eCAVIAR colocalisation

If for at least one CS in the overlap there was no log(BF) information (at least one CS is a result of PICS fine-mapping) we conducted colocalisation analysis with eCAVIAR[9], a heuristic algorithm that uses SNP PIPs. The colocalisation posterior probability (CLPP, equivalent to $H_4$ from the COLOC method) was computed as the sum over the product of overlapping variant fine-mapping probabilities between a GWAS and GWAS/QTL study.

##### 4.9. Calculation of colocalisation directionality

To determine effect directionality for each colocalising locus, the ratios between the two studies' betas are used. Within a locus, at each variant with a non-zero beta, the ratio between variant betas is calculated and the sign of these ratios (+1 or $-1$) indicates the variant's directionality. The average of these signs across a locus indicates the locus directionality. Where the average locus sign is 1 or $-1$ the directionality is the same or opposite, respectively. Where the locus is neither $-1$ nor 1, the directionality is inconclusive.

##### 4.10. Feature matrix generation and features description

For each gene assigned to a credible set, we assessed 28 functional genomic features and generated a feature matrix. Features are categorised into two groups: *individual gene features*—assessed independently for each gene—and *neighbourhood features*—individual gene features divided by the maximum value of the feature across the genes assigned to a credible set. Neighbourhood features reflect each gene's performance relative to other genes in the region.

The feature matrix was generated in three steps. In the first step, for each individual gene feature, we assigned genes to the credible set and scored the value for that feature. The rules for assigning genes to the CS differ per feature. All features (except 'number of genes') were constructed to be in the range $[0, 1]$, where 0 represents no evidence and 1 is the maximum possible positive value. In the second step, for each gene assigned to the CS, the corresponding neighbourhood feature was assessed:

$$\text{neighbourhood feature} = \frac{\text{individual gene feature}}{\max_{\text{genes in CS}} \text{individual gene feature}}.$$

If the maximum value was 0, the neighbourhood feature was also set to 0. In the third step, the long-format feature matrix is converted to wide format, with null values filled with zeros.

The individual gene features and corresponding neighbourhood features divide into three categories:

1. **Distance features**: the physical distance from credible set variants to the TSS or footprint of genes. Mean distance features are calculated as a weighted sum of distance scores across all variants in the CS weighted by PIP. Sentinel distance features are evaluated with respect to the lead variant in the CS.

2. **Molecular QTL colocalisation features**: includes both summary statistic-derived colocalisation evidence and LD-derived colocalisation evidence (e.g., eCAVIAR). For GWAS signals with multiple independent molecular trait signals, the maximum colocalisation score across estimates is used.

3. **Variant pathogenicity features**: reflect VEP-derived pathogenicity scores. VepMean is a PIP-weighted sum of VEP scores across all variants in the CS.

##### 4.11. Training and test sets for L2G model

The goal of the L2G model is to assign genes to GWAS loci based on an analysis of various genomic and functional features. To build the training and test datasets we needed to assign true positive and true negative status to the subset of feature matrix rows (credible set–gene pairs), following a three-step procedure.

**The effector gene list.** We created the effector gene list representing the set of biologically validated gene–trait associations. This list was derived by combining several sources: (1) manually curated "gold standards" from the previous Open Targets Genetics portal (release 22.10)[3] (medium and high confidence); (2) gene–EFO mappings containing drug IDs that have passed Phase 3 or Phase 4 clinical trials from the latest ChEMBL release; (3) gene–EFO mappings with a score $\geq$ 0.95 from ClinVar, UniProt, Gene2Phenotype, Genomics England PanelApp, ClinGen, UniProt Literature and Orphanet from the 25.06 Open Targets Platform release. The combined list was de-duplicated to ensure uniqueness of gene–EFO pairs.

**The positive and negative CS–gene pair assignment.** For each protein-coding gene–EFO pair from the effector gene list, we extracted all corresponding rows from the feature matrix. To form the final list of positives, we applied the following filtering procedure:

1. Selected only protein-coding genes.
2. Removed any positive pairs with a `distanceSentinelTSS` feature less than 0.
3. For each positive gene–EFO pair, selected only replicated credible sets.
4. Among all positive CS–gene pairs, removed duplications based on 11 columns: `geneId`, `efo_terms`, `variantId`, `vepMaximum`, `vepMean`, `eQtlColocClppMaximum`, `pQtlColocClppMaximum`, `sQtlColocClppMaximum`, `eQtlColocH4Maximum`, `pQtlColocH4Maximum`, `sQtlColocH4Maximum`. Numeric columns were rounded to the second digit before de-duplication.
5. Removed credible sets consisting of more than two positives.

To define the negative CS–gene pairs, we selected all other protein-coding genes in the window of each credible set that did not have a strong protein–protein interaction (StringDB score $\geq$ 0.8) with any of the positive genes.

The final dataset was split into training and held-out test sets based on unique positive genes (80%/20%, respectively).

##### 4.12. L2G model training

The L2G model is a binary classifier based on a gradient boosting algorithm implemented in the `xgboost` library.

We implemented a robust cross-validation strategy to rigorously evaluate the L2G model using the training and held-out test sets defined above. The training set is used to fit the model and for cross-validation; the held-out test set provides an unbiased assessment of the final model's performance on completely new gene–trait associations. The cross-validation process involves splitting the training set into five folds, providing a less biased estimate of the model's average precision across multiple testing sets. Our current setup allows parametrisation of variables such as the depth of the tree, learning rate, and regularisation values through the configuration file.

##### 4.13. L2G predictions and associations

We applied the trained L2G model to each row of the feature matrix and obtained L2G predictions for each gene–credible set pair. These predictions were then aggregated at the level of EFO mappings into direct and indirect associations, yielding L2G scores for EFO–gene pairs. In the Open Targets Platform, direct associations are calculated as the harmonic sum of L2G scores across all credible sets linking a given gene to the same disease; indirect associations are calculated similarly, but aggregated across common ancestral disease IDs in the EFO ontology. For the analyses in this paper, we used the maximum L2G score across credible sets rather than the harmonic sum.
